# Still waters run deep: Large scale genome rearrangements in the evolution of morphologically conservative Polyplacophora

**DOI:** 10.1101/2024.06.13.598811

**Authors:** Julia D. Sigwart, Yunlong Li, Zeyuan Chen, Katarzyna Vončina, Jin Sun

**Author notes:** Authors for correspondence: Jin Sun, Julia D. Sigwart, Yunlong Li **Email:** < >, < >, < >. these authors contributed equally to this work (Julia D. Sigwart, Yunlong Li).

## Abstract

**Background:** A major question in animal evolution is how genotypic and phenotypic changes are related, and whether ancient gene order is conserved in living clades. Chitons, the molluscan class Polyplacophora, retain a body plan and general morphology apparently little changed since the Palaeozoic. We present a comparative analysis of five reference quality genomes, including four *de novo* assemblies, covering all major chiton clades, and an updated phylogeny for the phylum.

**Results:** We constructed 20 ancient molluscan linkage groups (MLGs) that are relatively conserved in bivalve karyotypes, but subject to re-ordering, rearrangement, fusion, or partial duplication among chitons, varying even between congeneric species. The largest number of novel fusions is in the most plesiomorphic clade Lepidopleurida, and the chitonid *Liolophura japonica* has a partial genome duplication, extending the occurrence of large-scale gene duplication within Mollusca.

**Conclusions:** The extreme and dynamic genome rearrangements in this class stands in contrast to most other animals, demonstrating that chitons have overcome evolutionary constraints acting on other animal groups. The apparently conservative phenome of chitons belies rapid and extensive changes in genome.

## Introduction

The genomic mechanisms that enable or limit the evolution of major morphological changes remains one of the great questions of evolutionary biology. Living members of Mollusca represent the broadest morphological disparity of any animal phylum: mollusc body plans encompass squid, worms, living rocks and candy-coloured tree snails, as well as diverse intermediate and additional novel forms known from the fossil record. Early work on chromosome numbers in meiotic division used molluscs, and interest in understanding patterns in chromosome numbers across species also led to fundamental insights in animal polyploidy. Despite playing a key role in early advances, this important phylum lags in terms of quality and taxonomic coverage of whole genome sequence data. Reconstructing the evolution of genome architecture through deep time in diverse molluscs remains critical to understand genome evolution in animals. High quality genomic data for the deeply divergent, morphologically constrained chitons, would be expected to offer an opportunity to explore ancient genetic traits and evolutionary mechanisms preserved across the long span of animal evolution.

Prior work has reasonably assumed that rates of intra-chromosomal gene translocation are constant within major groups (Mackintosh et al., 2023). If this is true, syntenic rearrangement could be a clocklike indicator of divergence times. But rates of genomic rearrangement are not well known in molluscs, nor how they might vary across this vast clade, and higher rates of rearrangement confound reconstruction of ancestral states. In molluscs, an excellent fossil record allows an independent record of divergence times that will provide more insight into the variability (and utility) of rates of rearrangement as a measure of divergence time.

Chitons have long been considered the key taxon to understanding the ancestral molluscan body plan (Wanninger and Wollesen, 2019). Around 1000 living chiton species worldwide all possess an eight-part shell armour that has remained superficially unchanged with relatively little variation since the Carboniferous, over 300 Million years ago (Sigwart, 2017). Chitons are increasingly important for bio-inspired design, which will benefit from genomic tools to understand genetic control of their flexible armour and unique sensory systems (Ampuero et al., 2024; Varney et al., 2021). Chitons are generally conserved, yet the known species richness in chitons is higher than for the more morphologically and behaviourally diverse extant cephalopods. The adaptations that drive speciation in this group remain an open question.

One key issue is whether polyplacophoran molluscs are conservative *per se* or whether their diagnostic body plan adaptations are so distinctive and strange that this has overshadowed full understanding of additional adaptive traits: fully innervated shells, iron mineralised radula, living at almost all depths and latitudes of the world ocean. While the body plan of chitons has persisted for over 300 My, this is a template for remarkable adaptations that have only recently begun to be appreciated.

Chitons are grazers and most are not highly ecologically specialised (Sigwart and Schwabe, 2017). These animals mostly have separate sexes and all are broadcast spawners, are not migratory, cannot reasonably be subject to strong sexual selection, and many similar species co-occur in parapatric or sympatric radiations (Kelly and Eernisse, 2008). Previously published karyotype data shows closely related species with overlapping ranges can have different numbers of chromosomes, such as *Acanthochitona discrepans* (1n=8) and *A. crinita* (1n=9) in the North Atlantic (Certain, 1951). Chromosome rearrangement could be an attractive speculative explanation that could support maintaining species boundaries in this clade.

Here, we sequenced four new reference-quality genomes that cover the three taxonomic orders of living chitons: *Deshayesiella sirenkoi* (Lepidopleurida), *Callochiton septemvalvis* (Callochitonida), *Acanthochitona discrepans* and *A. rubrolineata* (Chitonida). Callochitonida is sister to Chitonida (Moles et al., 2021) and together these could be considered the Chitonida *sensu lato*. One aim here was to test the potential genomic differences separating these two orders within Chitonida s.l. We used these four new genomes together with previously published genomic data to reconstruct a phylogeny for the phylum. This allows us to confidently reconstruct ancestral chromosome arrangement of total-group Mollusca, and at different transitions within Polyplacophora.

## Results

We sequenced chromosome-level genomes of *Deshayesiella sirenkoi* (Lepidopleurida) from the Daikoku vent field, Western Pacific Ocean, *Callochiton septemvalvis* (Chitonida s.l.: Callochitonida) and *Acanthochitona discrepans* (Chitonida *sensu stricto*) both from the intertidal of Strangford Lough, N. Ireland, and the congener *A. rubrolineata* from the intertidal of Qingdao, China. These were sequenced with PacBio HiFi and scaffolded using Hi-C, resulting in high quality assemblies with over 97% BUSCO completeness (94% for *Callochiton septemvalvis*) and with numbers of chromosomes that differed among genera from 8-13 (Fig. 1, SI Appendix, Fig. S1, Table S1). These species show high levels of heterozygosity, ranging from almost 1% in *Deshayesiella* to 4.12% in *Callochiton* (Fig. 1, SI Appendix, Fig. S2).

**Fig 1.**
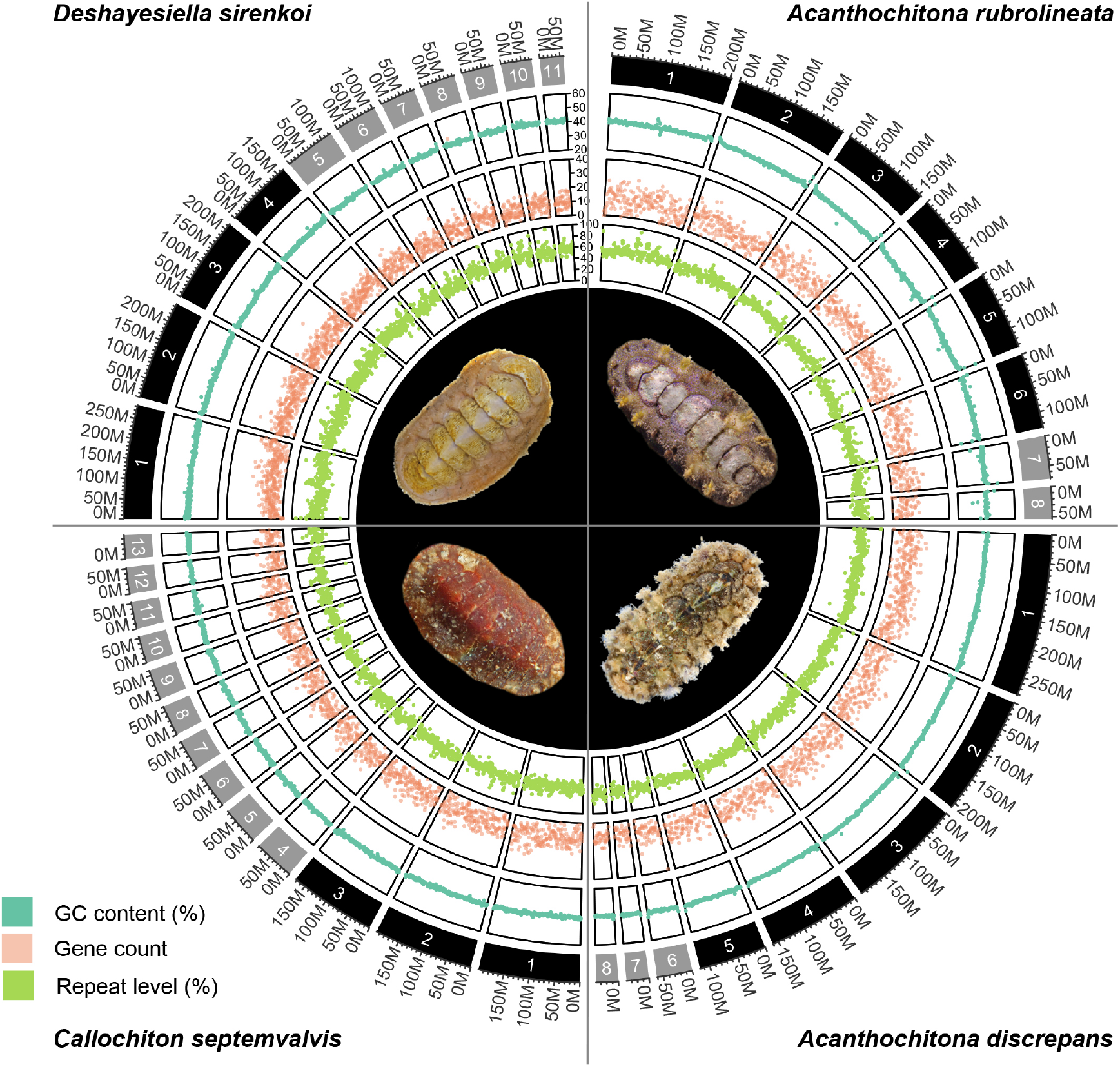
CIRCOS plots for 4 new chitons genome assemblies, clockwise from top left: one species in the order Lepidopleurida *Deshayesiella sirenkoi*, and three species in the clade Chitonida *sensu lato*: *Acanthochiton rubrolineata, A. discrepans*, and *Callochiton septemvalvis*. Each quarter circle shows the pseudochromosome content for each species, in order of size, with concentric rings indicating GC content, gene count, percent repeat content, and a photograph of the respective species.

Our phylogenetic results confirm the placement of chitons as sister to a monophyletic Conchifera, and we recover the expected topology within Polyplacophora consistent with other recent work using genomic and morphological characters (Fig. 2). Within Conchifera, we confirm the topology of other recent studies (Song et al., 2023); however, our supplementary analyses recovered Scaphopoda sister to Gastropoda but with lower support (SI Appendix, Fig. S3). Comparison with genomes from other molluscan classes shows the molluscan ancestor had a genome composed of 20 linkage groups (Fig. 2), we refer to these as the Molluscan Linkage Groups (MLG) 1-20. Three important fusion events are apparent synapomorphies for Polyplacophora, present in all living chitons and no Conchifera: MLG 4+16+18, MLG 7+10, and MLG 8+9 (Fig. 3, Fig. 4, SI Appendix, Figures S4-S7). There are additional fusions and intra-chromosomal rearrangements that are different in every species sampled.

**Fig 2.**
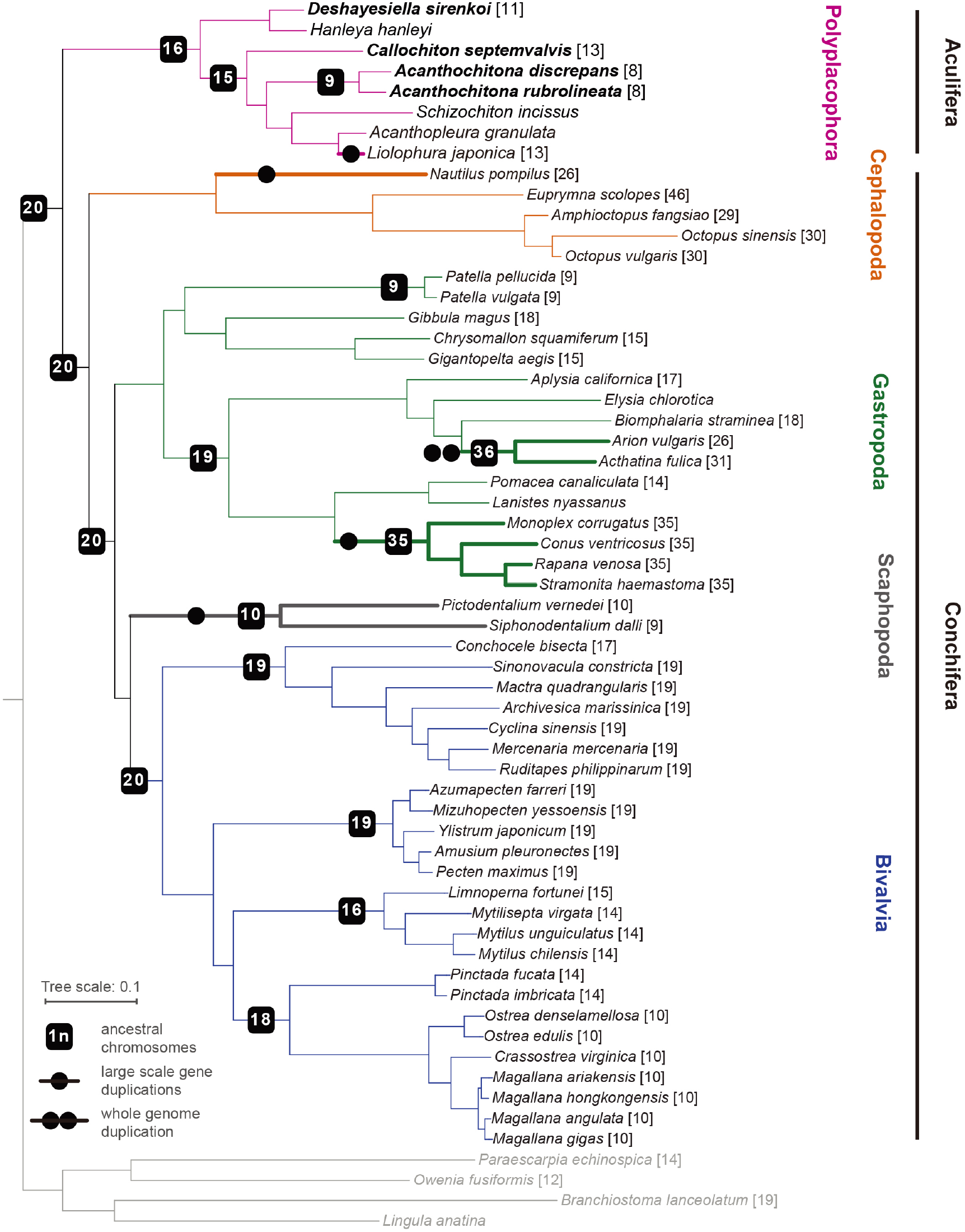
Phylogeny of Mollusca, with new genomes noted in bold type, and the chromosome number in square brackets for each species where known. Lineages with known whole (double circle) or partial genome (single circle) duplication are in thicker lines for emphasis; boxes on branches show the reconstructed ancestral 1n chromosome number for the respective clade.

**Fig 3.**
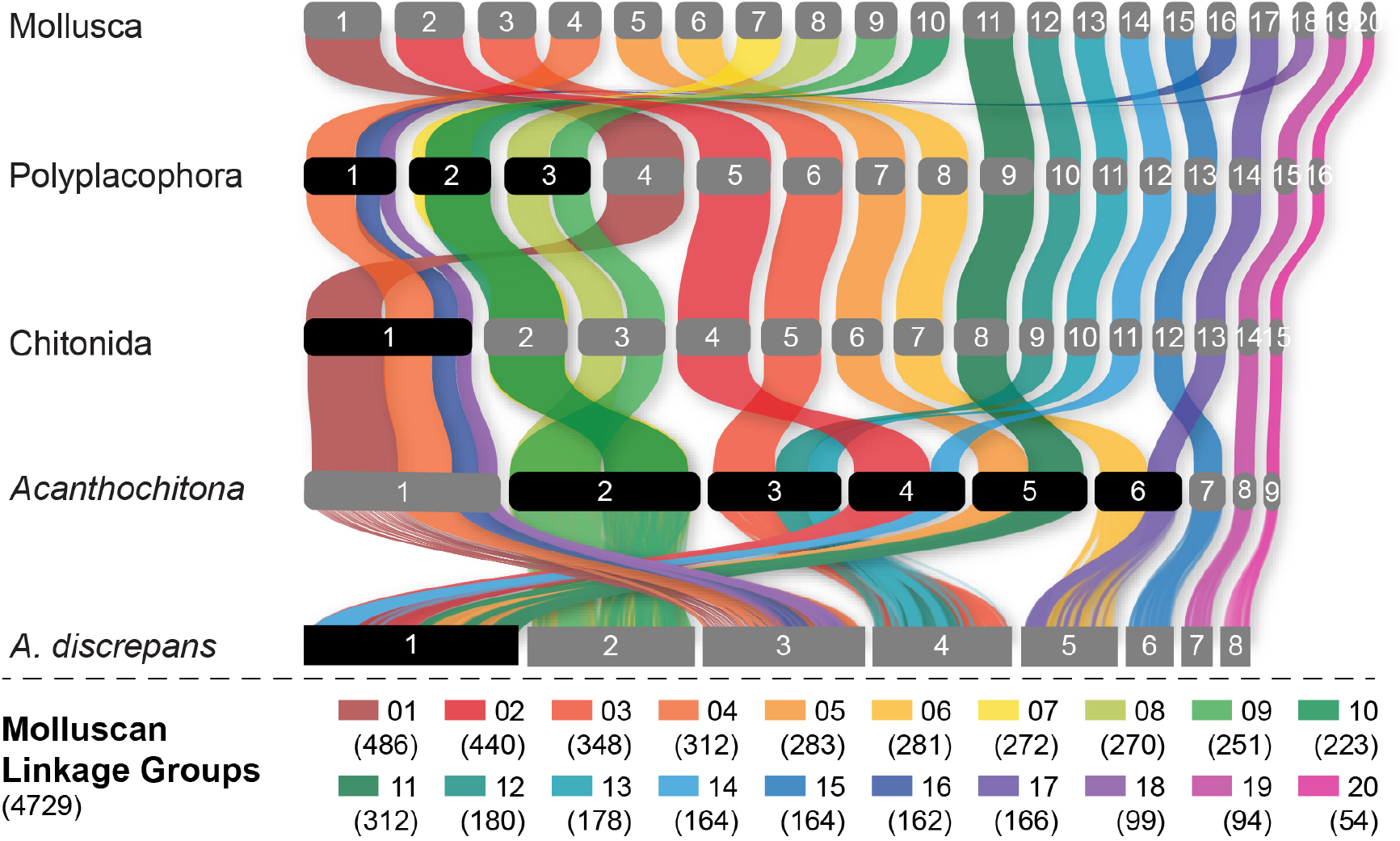
Evolution of the ancestral molluscan linkage groups (MLGs) within Polyplacophora using the lineage leading to *Acanthochitona discrepans* as an example. MLGs are distinguished by colours, at the top of the diagram and in the key at bottom showing the number of orthologs. Each row is the reconstructed karyotype of the ancestor of living Polyplacophora, the order Chitonida *sensu lato*, and the genus *Acanthochitona*. Reconstructed chromosomes on each row are numbered in order of size from largest (left) to smallest (right); chromosome fusions are highlighted with chromosome numbers in black boxes. This presentation highlights the extent of shifts especially in comparison to the molluscan or polyplacophoran ancestor.

**Fig 4.**
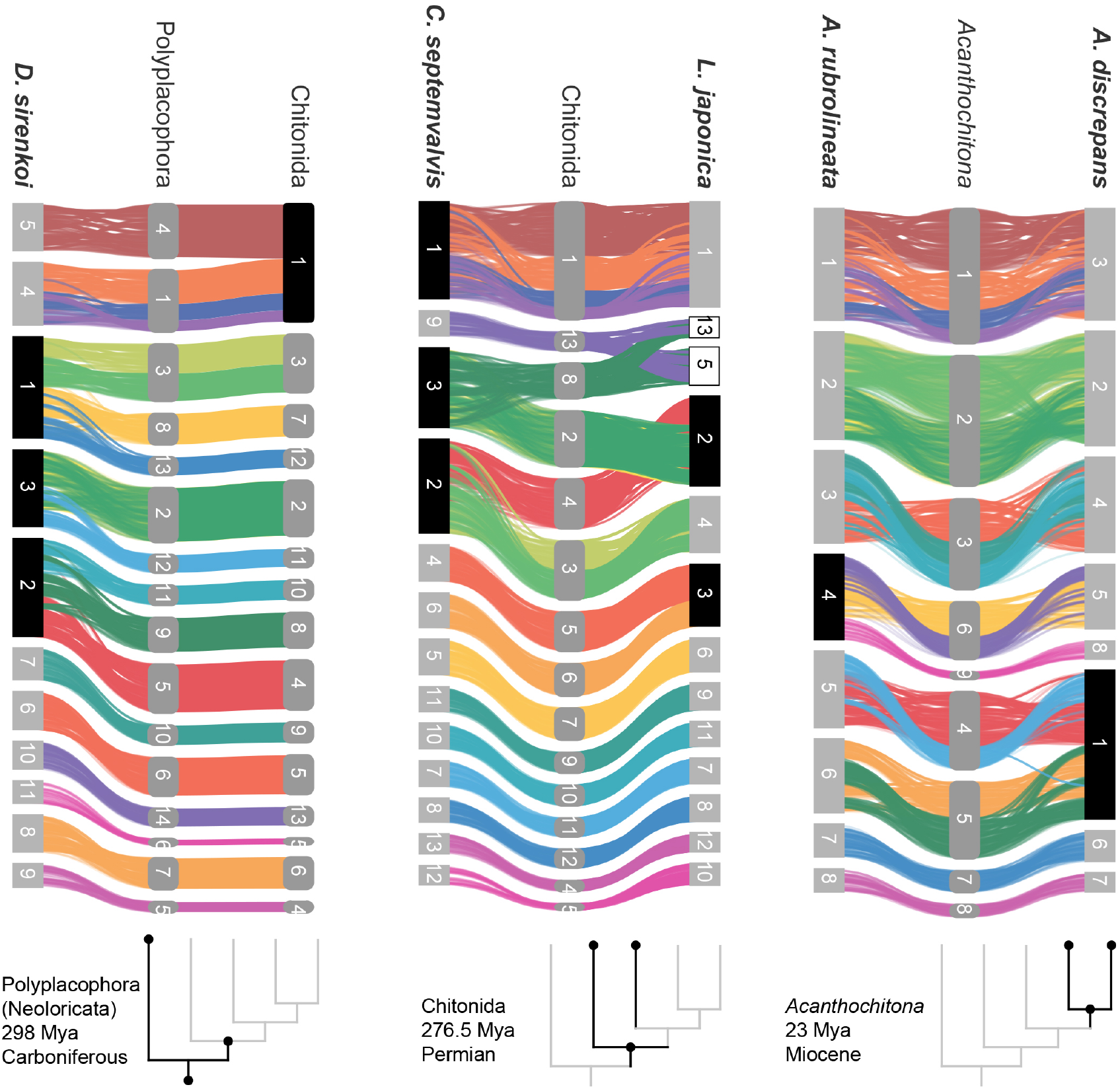
Syntenic rearrangements of MLGs within the evolution of Polyplacophora. Each part shows the reconstructed karyotypes of an ancestor (middle) and two descendent lineages, with a schematic cladogram for orientation. From left to right, the divergence of A) ancestor of Polyplacophora, leading to the lepidopleuran species *Deshayesiella sirenkoi* (left) and ancestor of Chitonida (right), B), ancestor of Chitonida s.l. leading to the callochitonid *Callochitona septemvalvis* (left) and chitonid *Liolophura japonica* (right) and C) ancestor of the genus *Acanthochitona* leading to the two congeneric species *A. rubrolineata* (left) and *A. discreapans* (right). Colours and presentation are as in Figure 3, and chromosome numbers indicate the sequence in terms of size from largest (1) to smallest. Here, the chromosomes are not in order of size but reordered such that each transition from the nearest ancestor is visible in more detail. Chromosome fusions are highlighted with chromosome numbers in black boxes, and duplications in white boxes.

## Discussion

### Chiton genomes show frequent and extreme rearrangement

Four new chiton genomes represent the most complete genomes sequenced for the class, adding to several previously published partial or complete genomes (Hong Kong Biodiversity Genomics Consortium et al., 2024; Liu et al., 2023; Varney et al., 2022, 2021) (SI Appendix, Tables S1-S3). Our comparisons of conchiferan and aculiferan (polyplacophoran) linkage groups confirm previous studies that also predicted a haploid karyotype of 20 for the molluscan ancestor based on other metazoans (Simakov et al., 2022).

Previous work demonstrated the variability in chiton karyotypes; species in the clade Chitonida have reported haploid numbers ranging maximally from 6 to 16 (Odierna et al., 2008) with a mode of 11 (Hallinan and Lindberg, 2011). Our data give the first indication for expected chromosome count in a species in Lepidopleurida (1n=11) within this established range. Using five chromosome-level genome assemblies for chitons, we reconstructed the ancestral karyotype for Polyplacophora (more strictly the taxonomic order Neoloricata), and all intermediate phylogenetic nodes to demonstrate the stepwise fusion and rearrangement of gene linkage groups during chiton evolution (Fig. 3).

Chitons demonstrate extreme genome rearrangement, even within a single genus. This represents not only gene order differences but syntenic (co-occurrence) changes in genomic architecture. Species of the genus *Acanthochitona* have a relatively short divergence time of maximally ∼23 My based on the fossil record (Dell’Angelo et al., 2020). *Acanthochitona discrepans* and *A. rubrolineata* each have 8 haploid chromosomes but these result from two different fusions compared to the reconstructed ancestral karyotype of *Acanthochitona* (Fig. 1, Fig. 4C). Previous karyotype data for *A. discrepans* were actually based on specimens of *A. crinita*; the species are very similar but the correct identification can be determined based on the geographic distributions (Vončina et al., 2023). *Acanthochitona crinita* has 9 haploid chromosomes (Certain, 1951): this is yet another syntenic arrangement within the same genus. There are major changes between congeners in different ocean basins (the Pacific *A. rubrolineata*) but also between two species in the NE Atlantic (*A. discrepans* and *A. crinita*) that are morphologically and ecologically almost indistinguishable.

Living species of Lepidopleurida retain more plesiomorphic morphology and this clade has a deep fossil record extending to the lower Carboniferous (Sigwart, 2017). The largest number of novel fusions among chiton genomes is three, in the lepidopleuran *Deshayesiella* (Fig. 4A). This in particular contrasts with the naive expectation that gene arrangement in most chiton genomes would be relatively conserved; Lepidopleurida retain plesiomorphic conditions in their morphology (Ampuero et al., 2024; Sigwart, 2017) yet this exemplar shows deviations from the reconstructed ancestral karyotype.

The remaining living chitons (Chitonida s.l.) comprise two sister clades recognised as separate orders: Chitonida and Callochitonida (Moles et al., 2021). *Callochiton* also has two additional fusion events, and the chitonid *Liolophura japonica* has a partial genome duplication, with two linkage groups fused apparently at first and then duplicated (Fig. 4B). In our reconstruction of ancestral karyotypes, there is no differences in arrangement between the ancestor of Chitonida *sensu stricto* or the ancestor of Chitonida *sensu lato* (Chitonida + Callochitonida). The four species in Chitonida s.l. share a large, fused chromosome (MLG 01+04+16+18) that is notably not well-mixed in *Liolophura japonica* (i.e., Chr01 in *Liolophura japonica*, Chr01 in *Callochiton septemvalvis*, Chr01 in *Acanthochitona rubrolineata*, and Chr03 in *A. discrepans*). Part of this pattern has its origin in the ancestral chiton karyotype and is retained in the lepidopleuran *Deshayesiella* (i.e., MLG 04+16+18). This implies a variable rate of intra-chromosomal rearrangement, with several of the MLGs conserved.

### Ancestral karyotypes for Mollusca

Chromosome numbers are not strongly conserved in animals. Changes in chromosome numbers are hypothesized to be an important driver of the diversification of Lepidoptera, with a strong correlation shown between rates of chromosome number changes and speciation (de Vos et al., 2020). The instability in lepidopteran chromosome numbers has mainly been studied from karyotype data but recently confirmed in genomic data (Chen et al., 2019), and changes within genera are known to encompass both neutral and adaptive evolution (Lucek, 2018; Vershinina and Lukhtanov, 2017). However, the level of rearrangement may be less than that in Polyplacophora.

Even within a single species, chromosome fusions are not uncommon, with Roberstonian translocations occurring in roughly 1 out of every 1000 live human births (Wilch and Morton, 2018). A previous study speculated that chromosome loss in related clades of chitons may be the result of Robertsonian translocation (Odierna et al., 2008). Although we still lack information on the telomere, this simple mechanism is not a sufficient explanation for several more complex intrachromosomal rearrangements (e.g. MLG 7+10: Fig. 4). Chromosome fusion does not present any immediate reproductive barrier, yet chromosome rearrangements between sister species can act as Dobzhansky-Muller incompatibilities and generate reproductive isolation (de Vos et al., 2020). Chitons are broadcast spawners, so such barriers may be speculatively advantageous, but so too are bivalves where these syntenic rearrangements are not found.

In order to compare syntenic changes within Polyplacophora to other mollusc clades, we re-analysed all available molluscan genomes in context of the newly identified MLGs. For example, the divergence of the bivalve clade Imparadentia (Lucinida + Venerida) is estimated in the Silurian, ca. 430 Mya (Crouch et al., 2021), or more than 150 My deeper than the divergence of crown group Polyplacophora, yet species within Imparadentia have almost no syntenic rearrangement (SI Appendix, Fig. S7, S8). Bivalves in Pteriomorphia, which have an even deeper origin in the lower Ordovician, ca. 485 Mya, show some rearrangement but members of two orders (Arcida and Pectinida) are highly similar, while two species in Ostreida that are intensively cultivated differ from these others and from each other (SI Appendix, Fig. S8).

Potential adaptive roles may be connected to contrasting mobility of different MLGs. The smallest two MLGs are the most conserved across the five chiton genomes; MLG20 is the only group that remains separate in all bivalve and chiton taxa. This region is also not duplicated in partial genome duplications in Neogastropoda, but is duplicated in whole genome duplication in two hermaphroditic terrestrial gastropods (*Achatina fulica* and *Arion vulgaris*) (Fig. 5, SI Appendix, Fig. S5).

**Fig 5.**
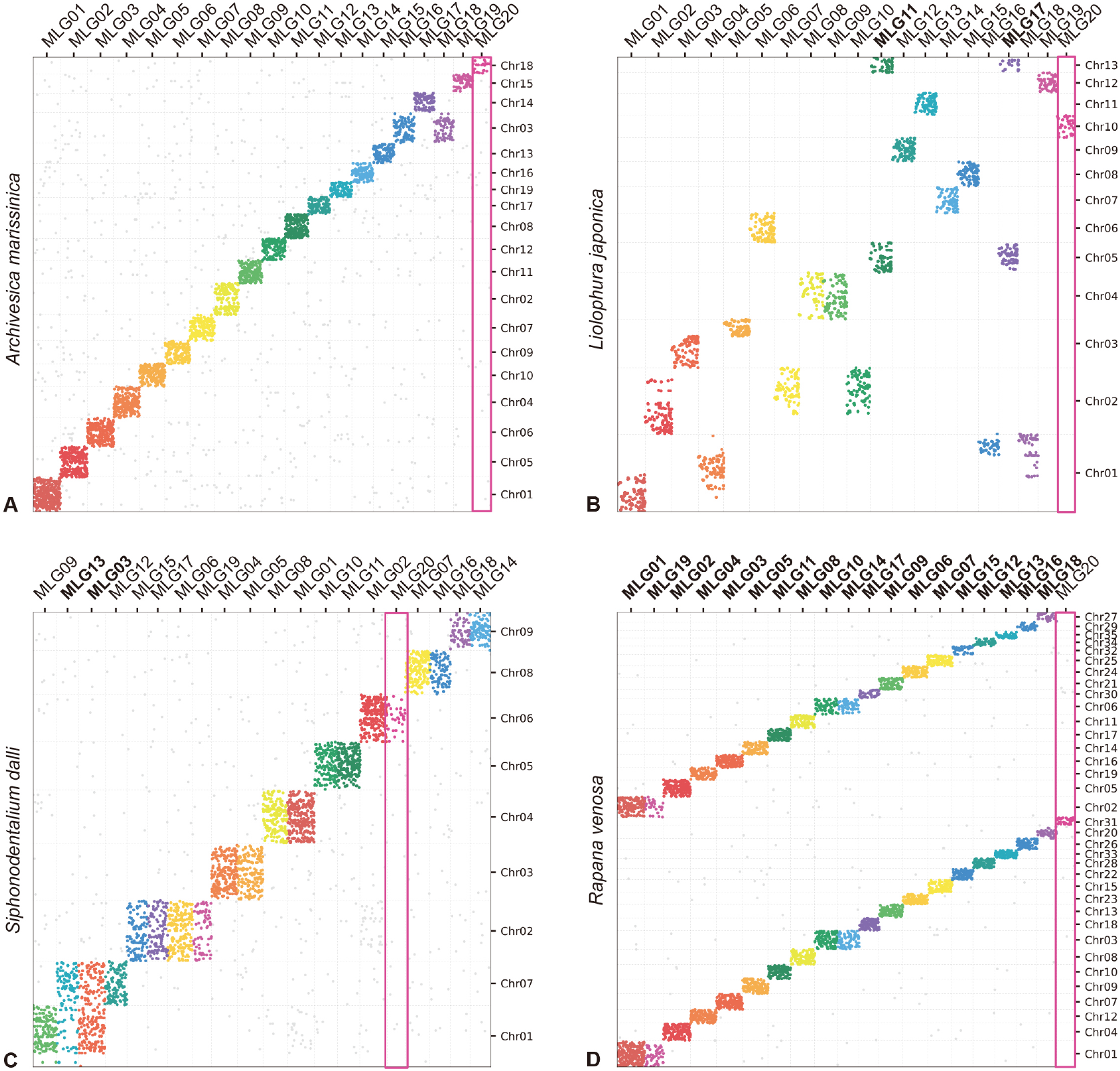
Oxford plots comparing gene occupancy of species from four different classes of molluscs (vertical) to the ancestral molluscan linkage groups (MLG, horizontal): A) the bivalve *Archivesica marisinica* retains a plesiomorphic karyotype reflecting the 20 MLGs, B) the chiton *Liolophura japonica* has large scale gene duplication in MLG11 and MLG17 on separate pseudochromosomes, C) the scaphopod *Siphonodetalium dalli* has large scale gene duplication of MLG13 and MLG03 on separate pseudochromosomes, D) the gastropod *Rapana venosa* demonstrates a nearly whole genome duplication with MLG20 not duplicated. Colours follow the presentation in Figure 3.

Earlier models based on karyotype data predicted three whole genome duplication events in the evolutionary history of living molluscs: in Neogastropoda, Stylommatophora, and coleoid cephalopods. Available genomes of Stylommatophora confirm whole genome duplication, as well as large scale gene duplications in Neogastropoda (SI Appendix, Fig. S6). In coleoid cephalopod genomes, the co-occurrence of loci from MLGs and intensive fusions (SI Appendix, Fig. S6) might not be easily resolved but may be more likely from the chromosome-disrupted processes reported in previous work (Albertin et al., 2023). By contrast, in *Nautilus* the linkage group that contains the conserved *hox* gene sets (Chr11, MLG16) shows no signal of co-occurrence with other MLGs or on other chromosomes (SI Appendix, Fig. S6). New results also show a partial and ancient genome duplication in the chiton *Liolophura* (Fig. 2, Fig. 4). Given that the MLGs are well conserved in all chitons if not their order (SI Appendix, Fig. S4), the event in *Liolophura japonica* is most likely a duplication instead of fission. Whole or partial genome duplication is now known from four molluscan classes, with contrasting patterns of ploidy (e.g. gastropods) or tandem (e.g. scaphopods) duplications (Fig. 5).

Reconstructing genome evolution is naturally more difficult for groups with high rates of intra-chromosomal rearrangement (Farré et al., 2019; Muffato et al., 2023). These patterns of co-occurrence of loci on the same chromosomes (synteny) should also persist for longer in evolutionary time, compared to faster rates of change in gene order (Damas et al., 2022; Simakov et al., 2022). Syntenic changes do not necessarily follow overall rates of translocation, which do not differ obviously among molluscs (SI Appendix, Table S4, S5). Confounding effects of rapid gene order or linkage group changes are not limited to issues of phylogenetic distance, when even closely related species have significant syntenic differences. Nonetheless a focus on linkage group arrangements is a promising direction for the many unresolved questions of deep molluscan phylogeny. Recent studies have championed the importance of synteny-based studies on genome evolution, as a basis to understand deep divergences and also the general mechanisms that underlie genomic architecture, fusion, fission, and translocation (Mackintosh et al., 2023). A general trend in insect and vertebrates is for increasing chromosome numbers in derived lineages. Fusion events may be more common in more recently derived chitons or connected to specific adaptations.

## Conclusions

The relatively small number of reference quality genomes for most molluscan groups is a temporary limitation to these analyses. High heterozygosity seems to be very common in molluscs, and is clearly a feature of chitons, which causes difficulty for high quality genome assembly. The heterozygosity for *Callochiton* at 4.12% far exceeds the 2.95% genomic heterozygosity reported as “one of the highest” for Lepidoptera (García-Berro et al., 2023). Large scale analyses based on genomics are equivocal about the drivers of genetic diversity, and further data from diverse clades must be included.

The concept of “diverse groups” is mostly based on perceived variability, typically manifested via morphological, ecological, or genetic differences. As one interesting comparison, Lepidoptera is commonly regarded as super-diverse, but also represent a recognisably constrained form, with variation in striking difference in colours, patterns and wing shapes. While lepidopteran variability is visually conspicuous, chitons are apparently equally variable, as demonstrated through their genetic rearrangement. Chitons, despite exceeding 1000 species in diverse ecological niches, with a wide range of morphological adaptations, are considered a “minor” and “neglected” clade. The group is taxonomically challenging – on the one hand, because of the supposed morphological stasis as a group, on the other, because of their high interspecific variability. Our comparative analyses suggest chromosomal-level changes are a pattern throughout much of chiton evolution, since chromosomal rearrangements are found when comparing congeneric species (*Acanthochitona* spp., Fig 4C) and also across orders (Fig 4A, 4B).

Chitons exhibit chromosome rearrangements at an almost unprecedented level. This is clear in the present study, the first comparative genomic study for Polyplacophora, compared to orders of magnitude more data for better studied groups. All this poses a more general question on how we define variability, how we perceive it, and also, if we actually understand it at all? Recognition of variability in the genome and phenome is crucial in understanding and re-evaluating unbiased measures of diversity in overlooked groups of organisms. The relationship revealed between the unstable arrangement of chiton genomes and species diversity provides new insights into potential mechanisms for speciation and broader diversification within Mollusca.

## Methods

Specimens were collected alive and flash frozen in liquid nitrogen (*A. rubrolineata*) or frozen at -80°C and stored at -80°C. High molecular weight genomic DNA was extracted from the foot of an individual specimen of each species, following the guidelines of the SDS method, and used for PacBio high fidelity (HiFi) sequencing and Hi-C library preparation. The PacBio Sequel II / IIe instrument was used for sequencing in CCS mode. For transcriptome sequencing, five separate tissues were dissected from the same specimen as DNA extraction, and preserved in RNAlater: foot (F), perinotum (P), radula sac (R), shell edge (S), and visceral mass (V).

Adapters in the reads were checked and removed using HiFiAdapterFilt version 2.0. The genome survey was finished using jellyfish v2.2.10 and Genomescope v2.0 (Ranallo-Benavidez et al., 2020), implemented 21- and 23-mer, which produced the estimated genome size and heterozygosity level for each species (SI Appendix, Fig. S2). The four genomes were assembled de novo based on qualified HiFi reads using hifiasm v0.16.1 (Cheng et al., 2021), with some additional curation steps for *C. septemvalvis* because of the high heterozygosity (SI Appendix). Potential contamination in the genome was detected and removed using blobtools v1.1.1 (Laetsch & Blaxter, 2017). Duplicated haplotigs and overlaps were removed for *D. sirenkoi, A. discrepans* and *A. rubrolineata* using Purge_Dups v1.2.6 (Guan et al., 2020).

The aligner STAR version 2.7.10a or STAR v2.7.3a was employed to map RNA-seq data into the working genome data (Dobin et al., 2012). The resulting alignment file served as an important support in prediction methods. *Ab initio* gene prediction was performed on the repeat-masked assembly in comparison with other published genomes (SI Appendix). Summary information on assembly, gene model, and annotation is provided in Table S2.

We identified putative orthologous sequences shared among genomes or transcriptomes for 58 molluscs and 4 additional metazoans to generate an alignment for phylogenetic analysis using VEHoP (Li et al., 2024). The alignments were removed if the overlap among them was less than 20 amino acids and there were less than 75% of taxa sampled, then, each alignment was used to construct “approximately maximum likelihood” tree using FastTree version 2.1.11 (Price et al., 2010). Phylogenetic relationships were investigated using IQ-TREE version 2.1.3 (Minh et al., 2020) with the “-MFP” model to compute the best-fit model of each partition and 1000 ultrabootstraps to test the topological support. To test the impact on conchiferan topology we ran a second analysis excluding Solemyida, the earliest diverging clade in Bivalvia (SI Appendix, Fig. S3).

The method for identifying ancient linkage groups followed previously published works. We resconstructed the linkages of the orthologues in molluscs, with the demonstrated commands, and conserved and representative proteins and their presumptive locations (https://github.com/ylify/MLGs). In detail, the ancient and conserved linkage groups were inferred from 7 chromosome-level mollusk assembles, including 1 gastropod (*Gibbula magus*), 1 bivalve (*Mizuhopecten yessonensis*), and the 5 chitons (*A. rubrolineata, A. discrepans, C. septemvalvis, D. sirenkoi*, and *Liolophura japonica*). The obtained gene set consisted of 4729 homologs, which could be detected in all 7 assembles and located in 20 linkage groups. The ancestral states of nodes were predicted to trace the karyotype evolutionary route in Polyplacophora, from the common ancestor of Mollusca to the genus *Acanthochitona*. Similarly, the nodes with more than two chromosome-level genomes within Mollusca were also investigated for the chromosome number of their common ancestor. We also compared the translocation rates across Mollusca, based on the non-syntenic rate of change divided by estimated divergence time. In this study, 25 genomes from four classes were selected to calculated the translocation rate at the inter-chromosome level, including 2 scaphopods, 5 chitons, 8 bivalves, and 10 gastropods.

## Declarations

## Acknowledgments

We thank Carola Greve, Damian Baranski, Alexander Ben Hamadou, and Charlotte Gerheim of the Translational Biodiversity Genomics (TBG) project based in the Senckenberg Research Institute and Museum, Frankfurt, for support with lab work and sequencing. We thank the staff of the Queen’s University Marine Laboratory, Portaferry, N Ireland, and Chong Chen (JAMSTEC) for support with field work and specimen collection. This is contribution number 28 of the Senckenberg Ocean Species Alliance. The sampling of *Deshayesiella sirenkoi* was carried out under the Marine Scientific Research permit number MSR U2022-047 from the United States government, since South Chamorro Seamount is within the Commonwealth of the Northern Mariana Islands.

## Availability of data and materials

The four chiton genomes projects have been deposited with the NCBI BioProject, with *Acanthochitona discrepans* in PRJNA1114954, *A. rubrolineata* in PRJNA1114370, *Callochiton septemvalvis* in PRJNA1114372, *Deshayesiella sirenkoi* in PRJNA1114373. In each project, the whole-genome sequencing data, Hi-C data, RNA-seq data have been affiliated. The genome assembly and gene-model predictions are deposited at figshare (10.6084/m9.figshare.27894189). The constructed Molluscan Linkage Groups (MLGs) are available, with the conserved and representative sequences and their predictive corresponding locations (https://github.com/ylify/MLGs/).

The commands used in this study have been deposited on GitHub (https://github.com/ylify/MLGs/), including the genome assembly, repeats and coding regions prediction, phylogenomic inference, and mutual best hits and ancient linkage groups detections.

## Author contributions

JDS and JS conceived the study and collected the samples. YL and ZC performed the analyses. JDS and KV led the biological interpretation of the results and review of comparative information from other studies. JDS drafted the manuscript. All authors contributed to interpreting the results and writing the manuscript.

## References

Albertin CB, Medina-Ruiz S, Mitros T, Schmidbaur H, Sanchez G, Wang ZY, Grimwood J, Rosenthal JJC, Ragsdale CW, Simakov O, Rokhsar DS. 2022 Genome and transcriptome mechanisms driving cephalopod evolution. Nat Commun 13, 2427. doi:10.1038/s41467-022-29748-w

Ampuero A, Vončina K, Parkinson DY, Sigwart JD. 2024. Aesthete pattern diversity in chiton clades (Mollusca: Polyplacophora): Balancing sensory structures and strength in valve architecture. J Morphol 285:e21784. doi:10.1002/jmor.21784

Certain P. 1951. Le caryotype d’Acanthochites discrepans Brown. Comptes Rendus Hebd Séances Académie Sci 233:435–437.

Chen W, Yang X, Tetreau G, Song X, Coutu C, Hegedus D, Blissard G, Fei Z, Wang P. 2019. A high-quality chromosome-level genome assembly of a generalist herbivore, Trichoplusia ni. Mol Ecol Resour 19:485–496. doi:10.1111/1755-0998.12966

Cheng H, Concepcion GT, Feng, X Zhang H, Li H. 2021. Haplotype-resolved de novo assembly using phased assembly graphs with hifiasm. Nat Methods 18, 170–175.

Crouch NMA, Edie SM, Collins KS, Bieler R, Jablonski D. 2021. Calibrating phylogenies assuming bifurcation or budding alters inferred macroevolutionary dynamics in a densely sampled phylogeny of bivalve families. Proc R Soc B Biol Sci 288:20212178. doi:10.1098/rspb.2021.2178

Damas J, Corbo M, Kim J, Turner-Maier J, Farré M, Larkin DM, Ryder OA, Steiner C, Houck ML, Hall S, Shiue L, Thomas S, Swale T, Daly M, Korlach J, Uliano-Silva M, Mazzoni CJ, Birren BW, Genereux DP, Johnson J, Lindblad-Toh K, Karlsson EK, Nweeia MT, Johnson RN, Zoonomia Consortium, Lewin HA. 2022. Evolution of the ancestral mammalian karyotype and syntenic regions. Proc Natl Acad Sci 119:e2209139119. doi:10.1073/pnas.2209139119

de Vos JM, Augustijnen H, Bätscher L, Lucek K. 2020. Speciation through chromosomal fusion and fission in Lepidoptera. Philos Trans R Soc B Biol Sci 375:20190539. doi:10.1098/rstb.2019.0539

Dell’Angelo B, Lesport J-F, Alain C, Sosso M. 2020. The Oligocene to Miocene chitons (Mollusca: Polyplacophora) of the Aquitaine Basin, soutwestern France, and Ligerian Basin, western France. Part 2: Lepidochitonidae, Tonicellidae, Acanthochitonidae, Cryptoplacidae and Additions to Part 1 56:1–58.

Dobin A, Davis CA, Schlesinger F, Drenkow J, Zaleski C, Jha S, Batut P, Chaisson M, Gingeras TR. 2012. STAR: ultrafast universal RNA-seq aligner. Bioinformatics 29, 15–21.

Guan D, McCarthy SA, Wood J, Howe K, Wang Y, Durbin R. 2020. Identifying and removing haplotypic duplication in primary genome assemblies. Bioinformatics 36, 2896–2898.

Farré M, Kim J, Proskuryakova AA, Zhang Y, Kulemzina AI, Li Q, Zhou Y, Xiong Y, Johnson JL, Perelman PL, Johnson WE, Warren WC, Kukekova AV, Zhang G, O’Brien SJ, Ryder OA, Graphodatsky AS, Ma J, Lewin HA, Larkin DM. 2019. Evolution of gene regulation in ruminants differs between evolutionary breakpoint regions and homologous synteny blocks. Genome Res 29:576–589. doi:10.1101/gr.239863.118

García-Berro A, Talla V, Vila R, Wai HK, Shipilina D, Chan KG, Pierce NE, Backström N, Talavera G. 2023. Migratory behaviour is positively associated with genetic diversity in butterflies. Mol Ecol 32:560–574. doi:10.1111/mec.16770

Hallinan NM, Lindberg DR. 2011. Comparative Analysis of Chromosome Counts Infers Three Paleopolyploidies in the Mollusca. Genome Biol Evol 3:1150–1163. doi:10.1093/gbe/evr087

Hong Kong Biodiversity Genomics Consortium, Hui JHL, Chan TF, Chan LL, Cheung SG, Cheang CC, Fang JK-H, Gaitan-Espitia JD, Lau SCK, Sung YH, Wong CKC, Yip KY-L, Wei Y, Au MFF, So WL, Nong W, Hui TY, Leung BKH, Williams GA. 2024. Chromosome-level genome assembly of the common chiton, Liolophura japonica (Lischke, 1873). Gigabyte 2024:1–14. doi:10.46471/gigabyte.123

Laetsch D, Blaxter M. 2017. BlobTools: Interrogation of genome assemblies. F1000 Research 6.

Kelly RP, Eernisse DJ. 2008. Reconstructing a radiation: the chiton genus Mopalia in the north Pacific. Invertebr Syst 22:17–28. doi:10.1071/IS06021

Li YL, Liu X, Cheng C, Qiu JW, Kocot KM, Sun J. 2024. VEHoP: A Versatile, Easy-to-use, and Homology-based Phylogenomic pipeline accommodating diverse sequences. bioRxiv 2024.07.24.604968; doi: 10.1101/2024.07.24.604968

Liu X, Sigwart JD, Sun J. 2023. Phylogenomic analyses shed light on the relationships of chiton superfamilies and shell-eye evolution. Mar Life Sci Technol 5:525–537. doi:10.1007/s42995-023-00207-9

Lucek K. 2018. Evolutionary Mechanisms of Varying Chromosome Numbers in the Radiation of Erebia Butterflies. Genes 9:166. doi:10.3390/genes9030166

Mackintosh A, Rosa PMG de la, Martin SH, Lohse K, Laetsch DR. 2023. Inferring inter-chromosomal rearrangements and ancestral linkage groups from synteny. doi:10.1101/2023.09.17.558111

Minh BQ, Schmidt HA, Chernomor O, Schrempf D, Woodhams MD, von Haeseler A, Lanfear R. 2020. IQ-TREE 2: New Models and Efficient Methods for Phylogenetic Inference in the Genomic Era. Mol Biol Evol 37, 1530–1534.

Moles J, Cunha TJ, Lemer S, Combosch DJ, Giribet G. 2021. Tightening the girdle: phylotranscriptomics of Polyplacophora. J Molluscan Stud 87:eyab019. doi:10.1093/mollus/eyab019

Muffato M, Louis A, Nguyen NTT, Lucas J, Berthelot C, Roest Crollius H. 2023. Reconstruction of hundreds of reference ancestral genomes across the eukaryotic kingdom. Nat Ecol Evol 7:355–366. doi:10.1038/s41559-022-01956-z

Odierna G, Aprea G, Barucca M, Biscotti MA, Canapa A, Capriglione T, Olmo E. 2008. Karyology of the Antarctic chiton Nuttallochiton mirandus (Thiele, 1906) (Mollusca: Polyplacophora) with some considerations on chromosome evolution in chitons. Chromosome Res 16:899–906. doi:10.1007/s10577-008-1247-1

Price MN, Dehal PS, Arkin AP. 2010. FastTree 2 – Approximately Maximum-Likelihood Trees for Large Alignments. PLOS ONE 5, e9490.

Ranallo-Benavidez TR, Jaron KS, Schatz MC. 2020. GenomeScope 2.0 and Smudgeplot for reference-free profiling of polyploid genomes. Nat Commun 11, 1432.

Sigwart JD. 2017. Deep trees: Woodfall biodiversity dynamics in present and past oceans. Deep Sea Res Part II Top Stud Oceanogr, Advances in deep-sea biology: biodiversity, ecosystem functioning and conservation 137:282–287. doi:10.1016/j.dsr2.2016.06.021

Sigwart JD, Schwabe E. 2017. Anatomy of the many feeding types in polyplacophoran molluscs. Invertebr Zool 14:205–216. doi:10.15298/invertzool.14.2.16

Simakov O, Bredeson J, Berkoff K, Marletaz F, Mitros T, Schultz DT, O’Connell BL, Dear P, Martinez DE, Steele RE, Green RE, David CN, Rokhsar DS. 2022. Deeply conserved synteny and the evolution of metazoan chromosomes. Sci Adv 8:eabi5884. doi:10.1126/sciadv.abi5884

Song H, Wang Yunan, Shao H, Li Z, Hu P, Yap-Chiongco MK, Shi P, Zhang T, Li C, Wang Yiguan, Ma P, Vinther J, Wang H, Kocot KM. 2023. Scaphopoda is the sister taxon to Bivalvia: Evidence of ancient incomplete lineage sorting. Proc Natl Acad Sci 120:e2302361120. doi:10.1073/pnas.2302361120

Varney RM, Speiser DI, McDougall C, Degnan BM, Kocot KM. 2021. The Iron-Responsive Genome of the Chiton Acanthopleura granulata. Genome Biol Evol 13:evaa263. doi:10.1093/gbe/evaa263

Varney RM, Yap-Chiongco MK, Mikkelsen NT, Kocot KM. 2022. Genome of the lepidopleurid chiton Hanleya hanleyi (Mollusca, Polyplacophora). doi:10.12688/f1000research.121706.1

Vershinina AO, Lukhtanov VA. 2017. Evolutionary mechanisms of runaway chromosome number change in Agrodiaetus butterflies. Sci Rep 7:8199. doi:10.1038/s41598-017-08525-6

Vončina K, Mikkelsen N, Morrow C, Ang R, Sigwart J. 2023. Clarification of the taxonomic status of Acanthochitona discrepans (Brown, 1827) with new data for the North-East Atlantic Acanthochitona (Polyplacophora, Acanthochitonidae). Biodivers Data J 11:e109554. doi:10.3897/BDJ.11.e109554

Wanninger A, Wollesen T. 2019. The evolution of molluscs. Biol Rev 94:102–115. doi:10.1111/brv.12439

Wilch ES, Morton CC. 2018. Historical and Clinical Perspectives on Chromosomal Translocations In: Zhang Y, editor. Chromosome Translocation. Singapore: Springer. pp. 1–14. doi:10.1007/978-981-13-0593-1_1

