## Supporting Information for "Still waters run deep: Large scale genome rearrangements in the evolution of morphologically conservative Polyplacophora"

Julia D. Sigwart *et al.*

**This PDF file contains**

**Appendix 1**. Extended Methods
**Supplementary References**
**Supplementary Figures** S1-S8
**Supplementary Tables** S1-S5

### Appendix 1. Extended Methods

*Sample collection, sequencing and assembly.*

In this work, four chiton species were collected for high quality genome assembly and annotation **(Table S1)**, including *Deshayesiella sirenkoi* (Lepidopleurida), *Callochiton septemvalvis* (Callochitonida), *Acanthochitona discrepans* and *A. rubrolineata* (Chitonida). Pathways for *A. discrepans* differ slightly as the complete sequencing and analysis was undertaken in Germany; pathways for *Callochiton septemvalvis* differ slightly because of problems encountered that we attribute to the highly heterozygous genome.

The foot was the target organ for genomic DNA extraction and the Hi-C library. The pooled strategy for RNA-seq was adopted, which included foot (F), perinotum (P), radula sac (R), shell edge (S), and visceral mass (V). The tissues were frozen at -80℃ (by liquid nitrogen for *A. rubrolineta*) and kept at -80℃ until needed.

Genomic DNA in the foot was extracted following the guidelines of the SDS method. PacBio high fidelity (HiFi) sequencing was adopted in genome assembly, with the platform and outputs shown in **Table S1**. The Ultra-Low-DNA-Input_Library was adopted in *C. septemvalvis* using SMRTbell Express Template Prep Kit 2.0 because of limited DNA yield for genomic sequencing. Genomic DNA of *C. septemvalvis* proceeded viaa next-generation sequencing in paired-end 150 bp mode, which was employed in a phasing step. Raw HiFi reads were extracted from the .bam files using extracthifi version 1.0.0 for *D. sirenkoi*, *C. septemvalvis*, *A. rubrolineata*, and DeepConsensus version 1.2.0 pipeline (Baid et al. 2022) for *A. discrepans*. Adapters in the reads were checked and removed using HiFiAdapterFilt version 2.0. The genome survey was finished using jellyfish v2.2.10 and Genomescope v2.0 (Ranallo-Benavidez et al., 2020), implemented 21- and 23-mer, which produced the estimated genome size and heterozygosity level for each sample **(Fig. S2**).

The qualified HiFi reads (i.e., adapter-free) were processed for *de novo* genome assembly using hifiasm v0.16.1-r375 (Cheng et al., 2021), which is proposed as one of the most time-efficient and optimal assemblers for HiFi reads. Regarding *C. septemvalvis*, the high heterozygosity level hindered the haploid genome assembly, so we adopted the phasing strategy. Specifically, we assembled the genome firstly using hifiasm with the disabled purge duplication (-l 0) and then obtained two pseudo-haploid genomes using khaper based on the feature of 17-mer (Zhang et al., 2021). Among them, the genome with better metrics (i.e., completeness and continuity) was selected as the representative genome of *C. septemvalvis*. For the other three chitons, the contig-level assemblies were obtained using hifiasm v0.16.1 (Cheng et al., 2021) with the default parameters. The potential contamination in the genome was detected and removed using blobtools v1.1.1 (Laetsch & Blaxter, 2017). Duplicated haplotigs and overlaps were removed for *D. sirenkoi*, *A. discrepans* and *A. rubrolineata* using Purge_Dups v1.2.6 (Guan et al., 2020).

For *D. sirenkoi*, *C. septemvalvis*, *A. rubrolineata*, the Hi-C reads were then aligned to the contig-level genome using the classical pipeline as demonstrated in previous work. In short, the valid reads (i.e., chimera from two primary DNA regions) were detected and extracted using HiC-Pro v3.1.0 (Servant et al., 2015), which integrated bowtie2 v2.5.1 (Langmead & Salzberg, 2012) and samtools v1.16.1 (Danecek et al., 2021) for reads mapping and check. They were further processed by the Juicer v1.6 (Durand et al., 2016) for the output of merged_nodups.txt and then the contigs were subjected to 3D de novo assembly (3D-DNA) pipeline v201008 (Dudchenko et al., 2017) with the default haploid setting for scaffolding. For *A. discrepans*, the Hi-C reads were aligned to the initial genome assembly using Arima Genomics mapping pipeline (https://github.com/ArimaGenomics/mapping_pipeline). In short, the reads were mapped to the reference using BWA-MEM v0.7.17-r1188 (Li 2013), converted to a sorted .bam file, and filtered to keep uniquely mapping pairs. PCR duplicates were removed using Picard v3.0.0 (Picard toolkit, 2019). The final alignment file and the assembly were then passed to YaHS v 1.1a-r3 (Zhou et al. 2023) for scaffolding in the default mode. Juicebox Assembly Tools (Dudchenko et al. 2018) were used to generate and visualize a Hi-C contact map. We manually curated the scaffolded assembly using an editable Hi-C heatmap to improve the assembly's quality and to correct misassembles with Juicebox v1.11.08 (Durand et al. 2016).

*Repeat region and gene models prediction*

The species-specific repeat element database in the contig-level haploid genome was *de novo* reconstructed using RepeatModeler v2.0.3 (Flynn et al., 2020), which was further adopted to identify and classify repeat regions in the genome using RepeatMasker v4.1.2-p1 (Smit et al., 2015). The identified repeat region in the genome was labeled as the lowercase bases, called a soft-masked genome.

The aligner STAR version 2.7.10a (Dobin et al., 2013) was employed to map RNA-seq data into the soft-masked genome. The .bam files were used in BRAKE2 version 2.1.6 (Brůna et al., 2021) which was implemented with Augustus version 3.4.0 and GeneMark version version 3.67_lic for the *ab inito* gene prediction, producing “augustus.gff3” and “genemark.gtf” as *ab-initio* predictions. These .bam files were also used in Trinity v2.13.2 and Stringtie v2.1.1 (Pertea et al., 2015) for transcripts assembly with genome-guided mode. Besides, the *de novo* assembled transcripts were obtained using Trinity, which could improve the completeness of transcripts. Both *de novo* assembled transcripts and genome-guided transcripts from Trinity were aligned to the soft-masked genome using PASA version 2.5.2 and aligner blat version 35, producing “pasa_aligned.gff3” as one of RNA-based predictions. Proteins from 28 high-quality genomes in metazoa were aligned to the soft-maked genome using miniport v0.5-r179 , producing “proteins.gff3” as the homology-based prodiction. Evidencemodeler v1.1.1 (Haas et al., 2008) was employed to merge or integrate the above three predictions into a comprehensive profile of genes, and the weights of evidence were AUGUSTUS for 2, GeneMark for 1, PASA for 8 or 10, Stringtie for 6, homolog from protein for 5. Genes were removed if they were solely supported by Augustus or GeneMark, due to the low confidence in the *ab-initio* predictions. The EVM consensus predictions were compared and updated with the de novo assembled transcripts using PASA v2.5.2 (Haas et al., 2008), which also led to the identification of untranslated regions (UTRs) and alternatively spliced isoforms in genes.

A slightly modified pipeline was employed in *A. discrepans*. Around 79M RNA-seq reads were aligned to the *A. discrepans* genome using STAR v2.7.3a (Dobin et al., 2013). The resulting alignment file served as an important support in all three prediction methods. *Ab initio* gene prediction was performed on the repeat-masked assembly with Braker3 (Gabriel et al. 2023) using default parameters. Two previously published chiton genomes, for *Acanthopleura granulata* (Varney et al. 2021) and *Hanleya hanleyi* (Varney et al. 2022) were selected and used for homology-based prediction. First, the proteins of *A. granulata* and *H. hanleyi* were downloaded from NCBI and aligned against the assembled genome using MMseqs2 (Hauser et al. 2016) with the parameter “-e 100.0 -s 8.5 --comp-bias-corr 0 --max-seqs 500 --mask 0 --orf-start-mode 1”. The results of homologous alignments were then combined into gene models with the splice site identified using mapped RNA-seq data with GeMoMa v1.9 (Keilwagen et al. 2019) using default parameters. Finally, the gene predictions were further sorted and filtered separately using the GeMoMa module GAF with default parameters. For the transcriptome-based prediction, the transcriptome of *A. discrepans* was assembled by both *de novo* and genome-guided approaches using Trinity v2.15.0 (Grabherr et al. 2011). The results were merged and passed to Program to Assemble Spliced Alignments (PASA) v2.5.2 (Haas et al., 2008) for gene predictions. In the end, all the predictions were combined into consensus coding sequence models using EVidenceModeler v1.1.1 (Haas et al., 2008), with the weighting of each method as “*ab initio* 1; homology-based 2, transcriptome-based 8”. In the end, we filtered out the incomplete genes and the predictions without any orthologs or transcripts supports using gFACs v1.1.2 (Caballero and Wegrzyn 2019).

The functional annotations of predicted protein sequences of chitons were obtained by searching public databases, including KEGG orthology by BlastKOALA (Kanehisa et al., 2016), gene ontology by blast2go version 5-basic against NCBI non-redundant protein database (nr). General information on assembly, gene model, and annotation is attached in **Table S2.**

*Phylogenomic relationships within Mollusca using high quality genomes*

The putative orthologous sequences shared among 62 metazoan genomes or transcriptomes (58 within Mollusca and 4 outside) were checked using OrthoFinder version 2.5.4 (Emms & Kelly, 2019), and then .fasta files in the ‘Orthogroup_Sequences’ directory were filtered under the pipeline modified from KM Kocot’ work (Kocot et al., 2017; Song et al., 2023). At first, .fasta files were removed unless at least 75% of taxa in a single file were sampled. The sequences in them were removed if the length was less than 100 amino acids or if they were redundant ones checked by uniqHaplo.pl. Mafft version 7.508 (Katoh & Standley, 2013) was applied to align sequences with the parameter of “--localpair --maxiterate 1000” and BMGE version 1.12 (Criscuolo & Gribaldo, 2010) was employed to trim ambiguously aligned columns in alignments. The alignments were removed if the overlap in them was less than 20 amino acids and there were less than 75% of taxa sampled, then, each alignment was used to construct “approximately maximum likelihood” tree using FastTree version 2.1.11 (Price et al., 2010) with the setting of “-slow -gamma,’ and then the paralogues in alignments were removed using phylopypruner version 1.2.6 (Thalen, 2018) with the setting of “--min-support 0.9 --mask pdist --trim-lb 3 --trim-divergent 0.75 --min-pdist 0.01 --prune LS”, which generated a supermatrix consisting of 4966 partitions. The phylogenetic relationship among them was investigated using IQ-TREE version 2.1.3 (Minh et al., 2020) with the “-MFP” model to compute the best-fit model of each partition and 1000 ultrabootstraps to test the topological support. In this work, Solemyida, the earliest divergent clade in Bivalvia, was removed for testing the position of Scaphopoda.

*Identification of ALGs and Conserved synteny in chromosome-level genomes*

The method for ancient linkage groups (ALGs) identification was reported in the two published works (Schultz et al., 2023; Simakov et al., 2022; Simakov et al., 2020). It should be noted that all the draft genomes for synteny were re-organized in the following ways: 1) the labels of chromosomes in a genome were re-ordered reversely according to their sizes, i.e., Chr01 equals to the longest chromosome; 2) these unanchored contigs/scaffolds and the corresponding genes in them were out of consideration; 3) the protein from the longest isoform was selected as the representative protein of gene if two or more isoforms in the gene. The identification of multi-way mutual best hit (MBH) clusters of encoding proteins among organisms or genomes is the core part. At first, the reciprocal blast hits between every two genomes were inferred by diamond blastp v2.1.8.162 (Buchfink et al., 2021) with the threshold of ‘-evalue 0.001 -max_target_seqs 50000’. Then, python was engaged in constructing the linkages of the orthologue among genomes (up to six), with the available scripts in Github (https://github.com/ylify/MLGs). Of which, only the strict one-to-one linkage was selected, meaning such an orthologue was presented in all genomes. For example, three proteins were shown as protein a in organism A, protein b in organism B, and protein c in organism C. The reciprocal blast hits revealed that 1) protein a and protein b, 2) protein b and protein c, 3) protein a and protein c, so a linkage of proteins a-b-c was defined.

The linkage groups between every two genomes were checked by Fisher’s exact test using R package macrosyntR (Sami El & Richard, 2023) with a significant threshold of p-value below 0.001, but it might define the group as insignificant one falsely if the number of linkages in the group was small. Therefore, the manual check of linkage groups between two genomes is needed using pairwise dot plots in JCVI version 1.2.7 (Tang et al., 2008), especially for organisms with genome duplication (e.g., *Acthatina fulica* and *Arion vulgaris*) or the recent insertion event (*Pomacea canaliculata*) or the two organisms with far relationships (e.g., Mollusca and Porifera). Coupling with the chromosomal location of proteins from .gff3 file, any potential linkage groups at the chromosome level would be constructed using ‘groups_six_species.py’ (a specified script). Accordingly, the ancient molluscan linkage groups (MLGs) were extracted based on the significant linkage groups on every two genomes. For example, similar to the definition of MBH, there were three chromosomal associations among three organisms: 1) chromosome 1 in organism A and chromosome 2 in organism B, 2) chromosome 1 in organism A and chromosome 3 in organism C, 3) chromosome 2 in organism B and chromosome 3 in organism C; then, it should be a linkage group from organism A-B-C as chromosome 1-2-3. Finally, the conserved collinearity among organisms was visualized as the vertical lines (linkages) and horizontal tracks (chromosomes in an organism) using JCVI version 1.2.7 (Tang et al., 2008). Colors in vertical lines were used to differentiate the linkage groups.

The ancestral states of nodes were predicted to trace the karyotype evolutionary route in Polyplacophora, from the common ancestor of Mollusca to the genus *Acanthochitona*. In detail, the ancient and conserved linkage groups were inferred from 7 chromosome-level mollusk assembles, including 1 snail (*Gibbula magus*), 1 bivalve (*Mizuhopecten yessonensis*), and 5 chitons (*A. rubrolineata*, *A. discrepans*, *C. septemvalvis*, *D. sirenkoi*, *Liolophura japonica*). The obtained gene set consisted of 4729 homologs, which could be detected in all 7 assembles and located in 20 linkage groups, with the name of mollusk linkage groups (MLGs). The AGORA v3.1 (Muffato et al., 2023) was employed to predict the gene order within linkage groups in ancestors, resulting in the assignment of genes in Contiguous Ancestral Regions (CARs). Then, the gene sets with the corresponding pseudo-locations in ancestral chromosomes were generated. We did not predict the sequences in the ancestral stats but integrated the proteins in the linkages (i.e., 7 proteins in a single linkage in this work). A customized Python script was used to check the chromosomal similarity between the MLGs and the extant genomes, with slight modifications from the identification of linkage groups. Specifically, diamond blastp was adopted to find the hits between them (MLGs as the query and proteins from genomes as databases). The protein was defined as a significant match against the mollusk ancestors if it was hit by at least 6 of 7 proteins from a linkage in MLGs (evalue < 0.001). Similarly, the significant linkage groups were checked by Fisher's exact test as mentioned before. Finally, the result was visualized as the Oxford plot using Matplotlib. The protein sequences and Python script are available on GitHub (https://github.com/ylify/MLGs). Similarly, the nodes with more than two chromosome-level genomes within Mollusca were also investigated for the chromosome number of their common ancestor. Moreover, the MLGs will help us to identify the ancestral stat and chromosomal events in Mollusca, including 1) the identification of chromosome duplication, fission, insertion, and fusion and 2) the real sense of whole genome duplication. To compare the syntenic changes between two genomes, the color scheme in MLGs was selected to visualize the classification of MBH. If the two proteins in MBH were both found in MLGs, the MBH would be defined as a significant linkage in Mollusca and then colored in oxford plot otherwise it would be in grey.

Translocation rates in species between chromosomes were evaluated as shown below; the non syntenic rate divided by the divergent time (based on calibrations from the fossil record). The non syntenic rate is the ratio of MBH not in a significant linkage group, compared to the total count of MBH. In this study, 25 genomes from four classes were selected to calculated the translocation rate at the inter chromosome level, including 2 scaphopods, 5 chitons, 8 bivalves, and 10 gastropods. The non syntenic rates of pairwise comparison were shown in **Table S4**. The divergent time among these four classes was set as 530 Mya according the first appearance of Bivalvia and Gastropoda (Benton et al., 2009; Bieler et al., 2014). The translocation rate of species was calculated by the mean value of non syntenic rate from the comparison against species from other classes. For example, the translocation rate in *A. discrepans* was the mean value of its comparison with the rest 20 non-chiton species. The results were shown in **Table S5**.

### References (Supplementary Materials)


Baid, G. et al. 2022. DeepConsensus improves the accuracy of sequences with a gap-aware sequence transformer. Nature Biotechnology.

Benton, M. J., Donoghuea, P. C. J., & Asher, R. J. (2009). Calibrating And Constraining Molecular Clocks. In S. B. Hedges & S. Kumar (Eds.), The Timetree of Life (pp. 0): Oxford University Press.

Bieler, R., Mikkelsen, P. M., Collins, T. M., Glover, E. A., González, V. L., Graf, D. L., . . . Girib et, G. (2014). Investigating the Bivalve Tree of Life – an exemplar-based approach combining molecular and novel morphological characters. Invertebrate Systematics, 28(1), 32-115. doi:https://doi.org/10.1071/IS13010

Brůna, T., Hoff, K. J., Lomsadze, A., Stanke, M., & Borodovsky, M. (2021). BRAKER2: automatic eukaryotic genome annotation with GeneMark-EP+ and AUGUSTUS supported by a protein database. NAR Genomics and Bioinformatics, 3(1). doi:10.1093/nargab/lqaa108

Buchfink, B., Reuter, K., & Drost, H.-G. (2021). Sensitive protein alignments at tree-of-life scale using DIAMOND. Nature Methods, 18(4), 366-368. doi:10.1038/s41592-021-01101-x

Caballero, M., and J. Wegrzyn. 2019. gFACs: Gene Filtering, Analysis, and Conversion to Unify Genome Annotations Across Alignment and Gene Prediction Frameworks. Genomics, Proteomics & Bioinformatics 17:305–310.

Cheng, H., Concepcion, G. T., Feng, X., Zhang, H., & Li, H. (2021). Haplotype-resolved de novo assembly using phased assembly graphs with hifiasm. Nature Methods, 18(2), 170-175. doi:10.1038/s41592-020-01056-5

Criscuolo, A., & Gribaldo, S. (2010). BMGE (Block Mapping and Gathering with Entropy): a new software for selection of phylogenetic informative regions from multiple sequence alignments. BMC Evolutionary Biology, 10(1), 210. doi:10.1186/1471-2148-10-210

Danecek, P., Bonfield, J. K., Liddle, J., Marshall, J., Ohan, V., Pollard, M. O., . . . Li, H. (2021). Twelve years of SAMtools and BCFtools. GigaScience, 10(2). doi:10.1093/gigascience/giab008

Dobin, A., Davis, C. A., Schlesinger, F., Drenkow, J., Zaleski, C., Jha, S., . . . Gingeras, T. R. (2013). STAR: ultrafast universal RNA-seq aligner. Bioinformatics, 29(1), 15-21. doi:10.1093/bioinformatics/bts635

Dudchenko, O., Batra, S. S., Omer, A. D., Nyquist, S. K., Hoeger, M., Durand, N. C., . . . Aiden, E. L. (2017). De novo assembly of the Aedes aegypti genome using Hi-C yields chromosome-length scaffolds. Science, 356(6333), 92-95. doi:doi:10.1126/science.aal3327

Durand, N. C., Shamim, M. S., Machol, I., Rao, S. S. P., Huntley, M. H., Lander, E. S., & Aiden, E. L. (2016). Juicer Provides a One-Click System for Analyzing Loop-Resolution Hi-C Experiments. Cell Systems, 3(1), 95-98. doi:https://doi.org/10.1016/j.cels.2016.07.002

Emms, D. M., & Kelly, S. (2019). OrthoFinder: phylogenetic orthology inference for comparative genomics. Genome Biology, 20(1), 238. doi:10.1186/s13059-019-1832-y

Flynn, J. M., Hubley, R., Goubert, C., Rosen, J., Clark, A. G., Feschotte, C., & Smit, A. F. (2020). RepeatModeler2 for automated genomic discovery of transposable element families. Proceedings of the National Academy of Sciences, 117(17), 9451-9457. doi:doi:10.1073/pnas.1921046117

Gabriel, L. et al. 2023. BRAKER3: Fully automated genome annotation using RNA-seq and protein evidence with GeneMark-ETP, AUGUSTUS and TSEBRA. <http://biorxiv.org/lookup/doi/10.1101/2023.06.10.544449> (11 June 2024).

Guan, D., McCarthy, S. A., Wood, J., Howe, K., Wang, Y., & Durbin, R. (2020). Identifying and removing haplotypic duplication in primary genome assemblies. Bioinformatics, 36(9), 2896-2898. doi:10.1093/bioinformatics/btaa025

Grabherr, M. G. et al. 2011. Full-length transcriptome assembly from RNA-Seq data without a reference genome. Nature Biotechnology 29:644–652.

Haas, B. J., Salzberg, S. L., Zhu, W., Pertea, M., Allen, J. E., Orvis, J., . . . Wortman, J. R. (2008). Automated eukaryotic gene structure annotation using EVidenceModeler and the Program to Assemble Spliced Alignments. Genome Biology, 9(1), R7. doi:10.1186/gb-2008-9-1-r7

Hauser, M., M. Steinegger, and J. Söding. 2016. MMseqs software suite for fast and deep clustering and searching of large protein sequence sets. Bioinformatics 32:1323–1330.

Kanehisa, M., Sato, Y., & Morishima, K. (2016). BlastKOALA and GhostKOALA: KEGG Tools for Functional Characterization of Genome and Metagenome Sequences. Journal of Molecular Biology, 428(4), 726-731. doi:10.1016/j.jmb.2015.11.006

Katoh, K., & Standley, D. M. (2013). MAFFT Multiple Sequence Alignment Software Version 7: Improvements in Performance and Usability. Molecular Biology and Evolution, 30(4), 772-780. doi:10.1093/molbev/mst010

Keilwagen, J., F. Hartung, and J. Grau. 2019. GeMoMa: Homology-Based Gene Prediction Utilizing Intron Position Conservation and RNA-seq Data. Pp. 161–177 in Gene Prediction (M. Kollmar, ed.). Springer New York, New York, NY.

Kocot, K. M., Struck, T. H., Merkel, J., Waits, D. S., Todt, C., Brannock, P. M., . . . Halanych, K. M. (2017). Phylogenomics of Lophotrochozoa with Consideration of Systematic Error. Systematic Biology, 66(2), 256-282. doi:10.1093/sysbio/syw079

Laetsch, D., & Blaxter, M. (2017). BlobTools: Interrogation of genome assemblies. F1000Research, 6(1287). doi:10.12688/f1000research.12232.1

Langmead, B., & Salzberg, S. L. (2012). Fast gapped-read alignment with Bowtie 2. Nature Methods, 9(4), 357-359. doi:10.1038/nmeth.1923

Li, H. 2013. Aligning sequence reads, clone sequences and assembly contigs with BWA-MEM. arXiv. <http://arxiv.org/abs/1303.3997> (11 June 2024).

Minh, B. Q., Schmidt, H. A., Chernomor, O., Schrempf, D., Woodhams, M. D., von Haeseler, A., & Lanfear, R. (2020). IQ-TREE 2: New Models and Efficient Methods for Phylogenetic Inference in the Genomic Era. Molecular Biology and Evolution, 37(5), 1530-1534. doi:10.1093/molbev/msaa015

Muffato, M., Louis, A., Nguyen, N. T. T., Lucas, J., Berthelot, C., & Roest Crollius, H. (2023). Reconstruction of hundreds of reference ancestral genomes across the eukaryotic kingdom. Nature Ecology & Evolution, 7(3), 355-366. doi:10.1038/s41559-022-01956-z

Pertea, M., Pertea, G. M., Antonescu, C. M., Chang, T.-C., Mendell, J. T., & Salzberg, S. L. (2015). StringTie enables improved reconstruction of a transcriptome from RNA-seq reads. Nature Biotechnology, 33(3), 290-295. doi:10.1038/nbt.3122

Price, M. N., Dehal, P. S., & Arkin, A. P. (2010). FastTree 2 – Approximately Maximum-Likelihood Trees for Large Alignments. PLOS ONE, 5(3), e9490. doi:10.1371/journal.pone.0009490

Ranallo-Benavidez, T. R., Jaron, K. S., & Schatz, M. C. (2020). GenomeScope 2.0 and Smudgeplot for reference-free profiling of polyploid genomes. Nature Communications, 11(1), 1432. doi:10.1038/s41467-020-14998-3

Sami El, H., & Richard, R. C. (2023). macrosyntR : Drawing automatically ordered Oxford Grids from standard genomic files in R. bioRxiv, 2023.2001.2026.525673. doi:10.1101/2023.01.26.525673

Schultz, D. T., Haddock, S. H. D., Bredeson, J. V., Green, R. E., Simakov, O., & Rokhsar, D. S. (2023). Ancient gene linkages support ctenophores as sister to other animals. Nature, 618(7963), 110-117. doi:10.1038/s41586-023-05936-6

Servant, N., Varoquaux, N., Lajoie, B. R., Viara, E., Chen, C.-J., Vert, J.-P., . . . Barillot, E. (2015). HiC-Pro: an optimized and flexible pipeline for Hi-C data processing. Genome Biology, 16(1), 259. doi:10.1186/s13059-015-0831-x

Simakov, O., Bredeson, J., Berkoff, K., Marletaz, F., Mitros, T., Schultz, D. T., . . . Rokhsar, D. S. (2022). Deeply conserved synteny and the evolution of metazoan chromosomes. Science Advances, 8(5), eabi5884. doi:doi:10.1126/sciadv.abi5884

Simakov, O., Marlétaz, F., Yue, J.-X., O'Connell, B., Jenkins, J., Brandt, A., . . . Rokhsar, D. S. (2020). Deeply conserved synteny resolves early events in vertebrate evolution. Nature Ecology & Evolution, 4(6), 820-830. doi:10.1038/s41559-020-1156-z

Smit, A., Hubley, R., & Green, P. (2015). RepeatMasker Open-4.0. 2013–2015. In.

Song, H., Wang, Y., Shao, H., Li, Z., Hu, P., Yap-Chiongco, M. K., . . . Kocot, K. M. (2023). Scaphopoda is the sister taxon to Bivalvia: Evidence of ancient incomplete lineage sorting. Proceedings of the National Academy of Sciences, 120(40), e2302361120. doi:doi:10.1073/pnas.2302361120

Tang, H., Bowers, J. E., Wang, X., Ming, R., Alam, M., & Paterson, A. H. (2008). Synteny and Collinearity in Plant Genomes. Science, 320(5875), 486-488. doi:doi:10.1126/science.1153917

Thalen, F. (2018). PhyloPyPruner: Tree-based Orthology Inference for Phylogenomics with New Methods for Identifying and Excluding Contamination.

Varney, R. M., D. I. Speiser, C. McDougall, B. M. Degnan, and K. M. Kocot. 2021. The Iron-Responsive Genome of the Chiton Acanthopleura granulata. Genome Biology and Evolution 13:evaa263.

Varney, R. M., M. K. Yap-Chiongco, N. T. Mikkelsen, and K. M. Kocot. 2022. Genome of the lepidopleurid chiton Hanleya hanleyi (Mollusca, Polyplacophora). F1000Research 11:555.

Zhang, X., Chen, S., Shi, L., Gong, D., Zhang, S., Zhao, Q., . . . You, M. (2021). Haplotype-resolved genome assembly provides insights into evolutionary history of the tea plant Camellia sinensis. Nature Genetics, 53(8), 1250-1259. doi:10.1038/s41588-021-00895-y

Zhou, C., S. A. McCarthy, and R. Durbin. 2023. YaHS: yet another Hi-C scaffolding tool. Bioinformatics 39:btac808.

**Supplementary Figures – Figure captions**

**Figure S1.** Hi-C contact map in pseudo-chromosome level assemblies of four chitons.

**Figure S2**. Genome survey of four chitons from the GenomeScope2 software.

**Figure S3.** Phylogenomic relationships among five classes in Mollusca (with two Solemya from RNA-seq). A single star in the tree indicates partial duplications and two stars indicate true whole genome duplication. The numbers in parentheses after taxon names indicate the number of pseudo-chromosomes where known, and the numbers on branches indicate the predicted (1n) number of ancestral chromosomes. The newly sequenced chiton species are indicated in bold text. In contrast to the tree presented in the main text, this tree resolved Scaphapoda + Gastropoda. IQTREE2: -m MFP -B 1000 (60 genomes and 2 transcriptomes)

**Figure S4.** Oxford plots of 5 chitons against MLGs

**Figure S5.** Oxford plots of 12 gastropods against MLGs, with the identification of the true sense of whole genome duplications (WGD). MLG20 is highlighted with a red box to distinguish its dynamics under WGD.

**Figure S6.** Oxford plots of 4 cephalopods and 2 scaphopods against MLGs.

**Figure S7.** Oxford plots of 6 bivalves against MLGs.

**Figure S8.** Syntenic graph in Bivalvia, highly conserved synteny in Imparadentia and large rearrangements within Peteriomorphia. Two clades in Bivalvia have 19 pseudo-chromosomes from a single fusion, but they are independent occurrences: in Pectinida, the fusion is MLG12 and MLG16, whereas in Venerida, the fusion is MLG16 and MLG18.

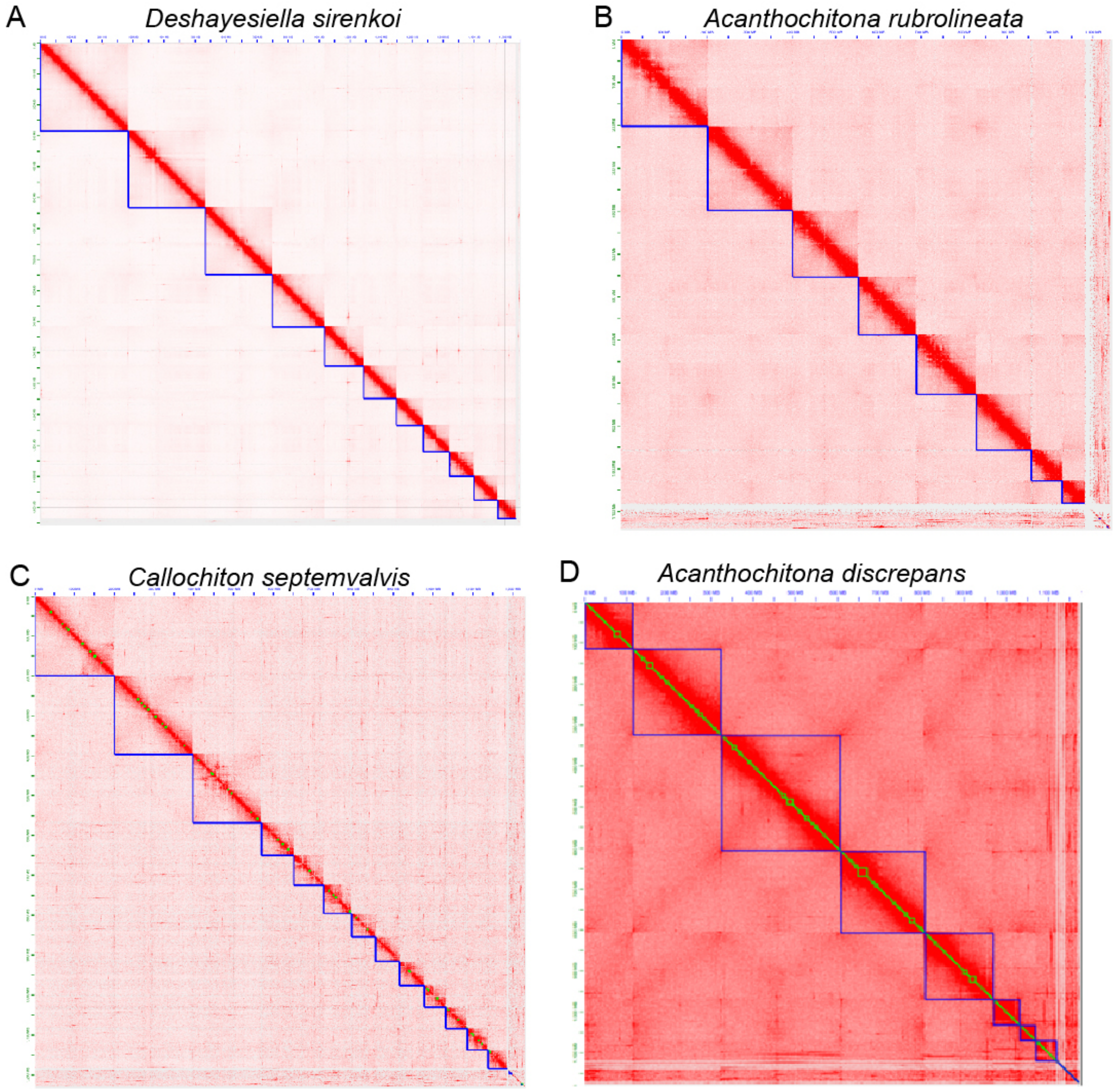


**Figure S1.** Hi-C contact map in pseudo-chromosome level assemblies of four chitons.

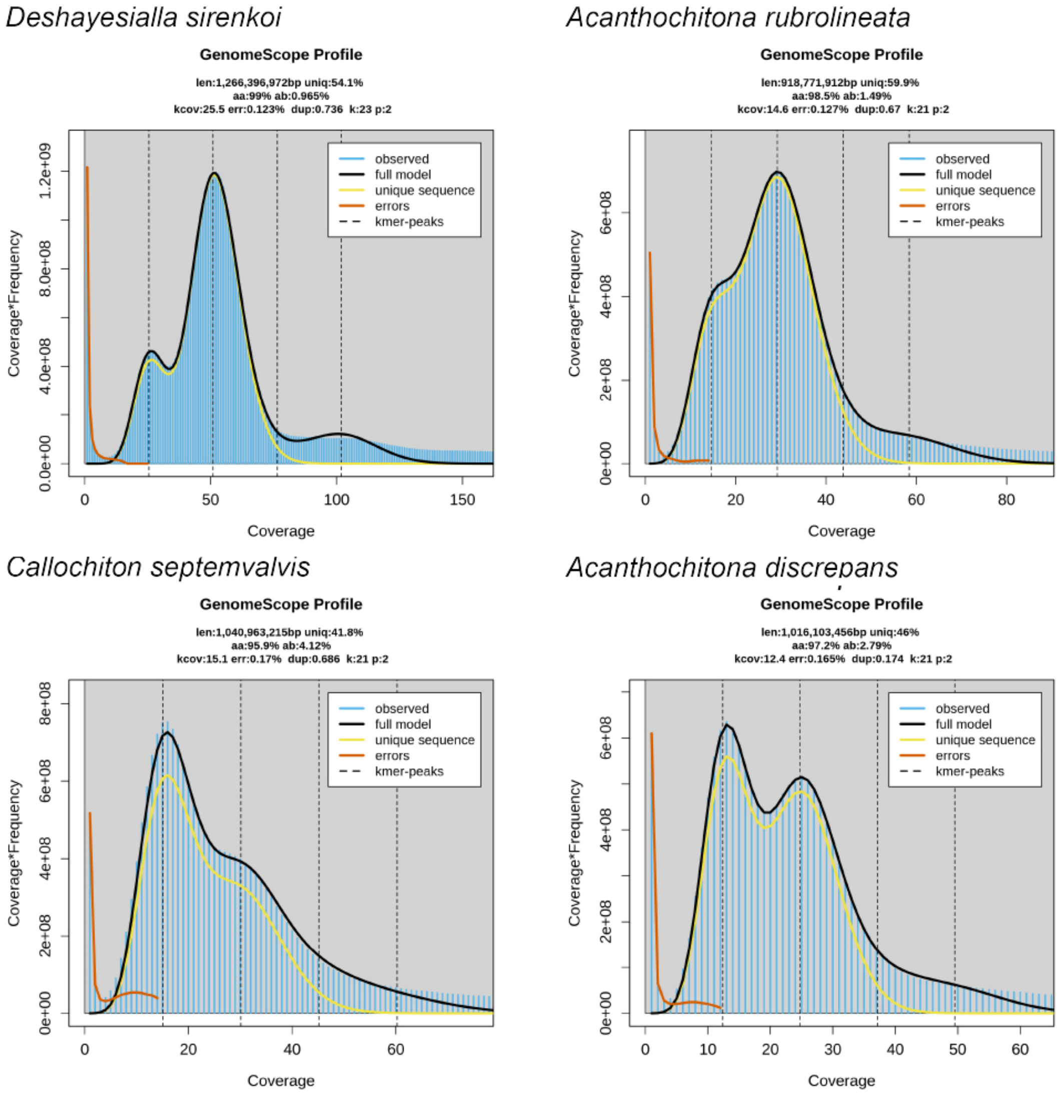


**Figure S2**. Genome survey of four chitons from the GenomeScope2 software.

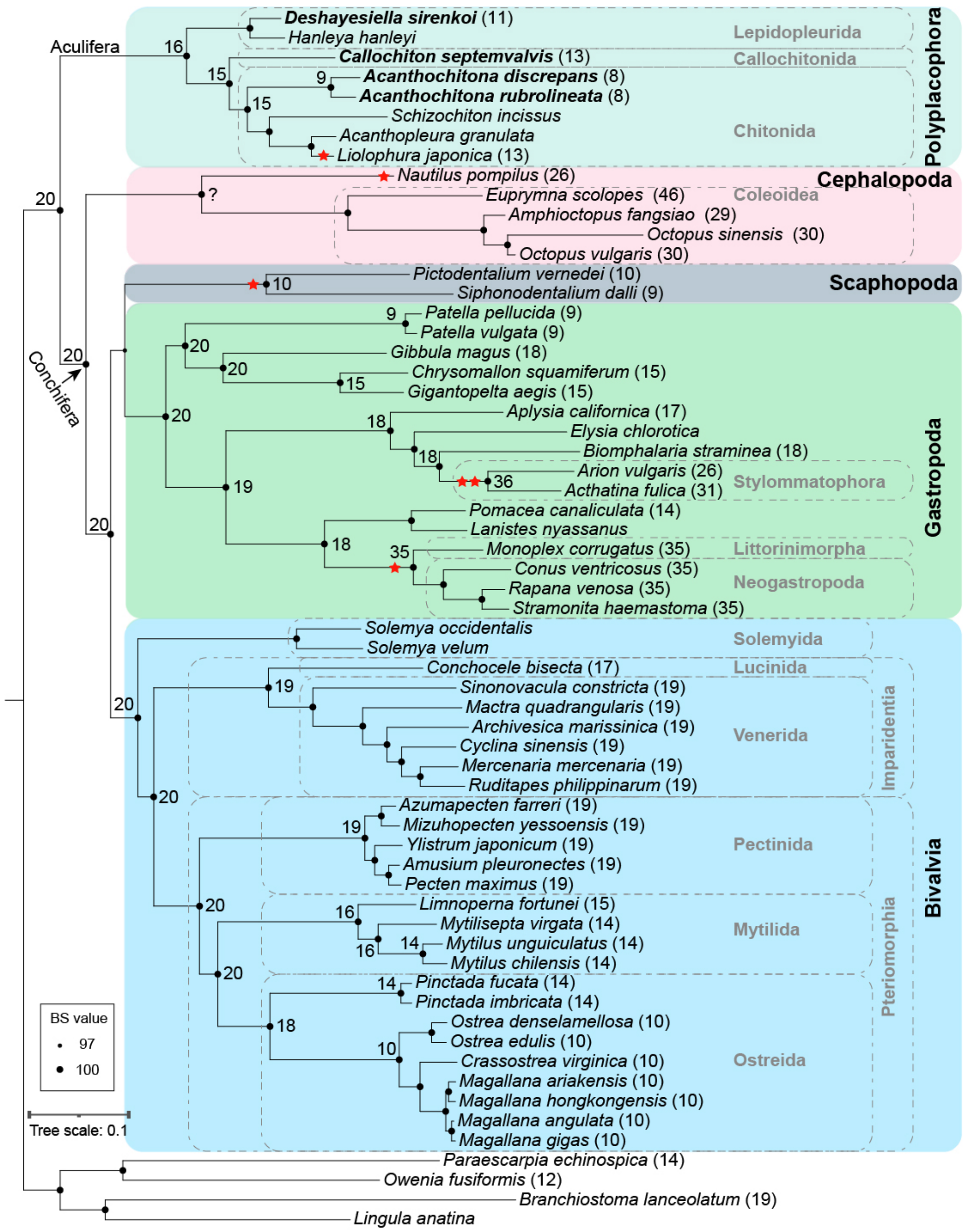


**Figure S3.** Phylogenomic relationships among five classes in Mollusca (with two Solemya from RNA-seq). A single star in the tree indicates partial duplications and two stars indicate true whole genome duplication. The numbers in parentheses after taxon names indicate the number of pseudo-chromosomes where known, and the numbers on branches indicate the predicted (1n) number of ancestral chromosomes. The newly sequenced chiton species are indicated in bold text. In contrast to the tree presented in the main text, this tree resolved Scaphapoda + Gastropoda. IQTREE2: -m MFP -B 1000 (60 genomes and 2 transcriptomes)

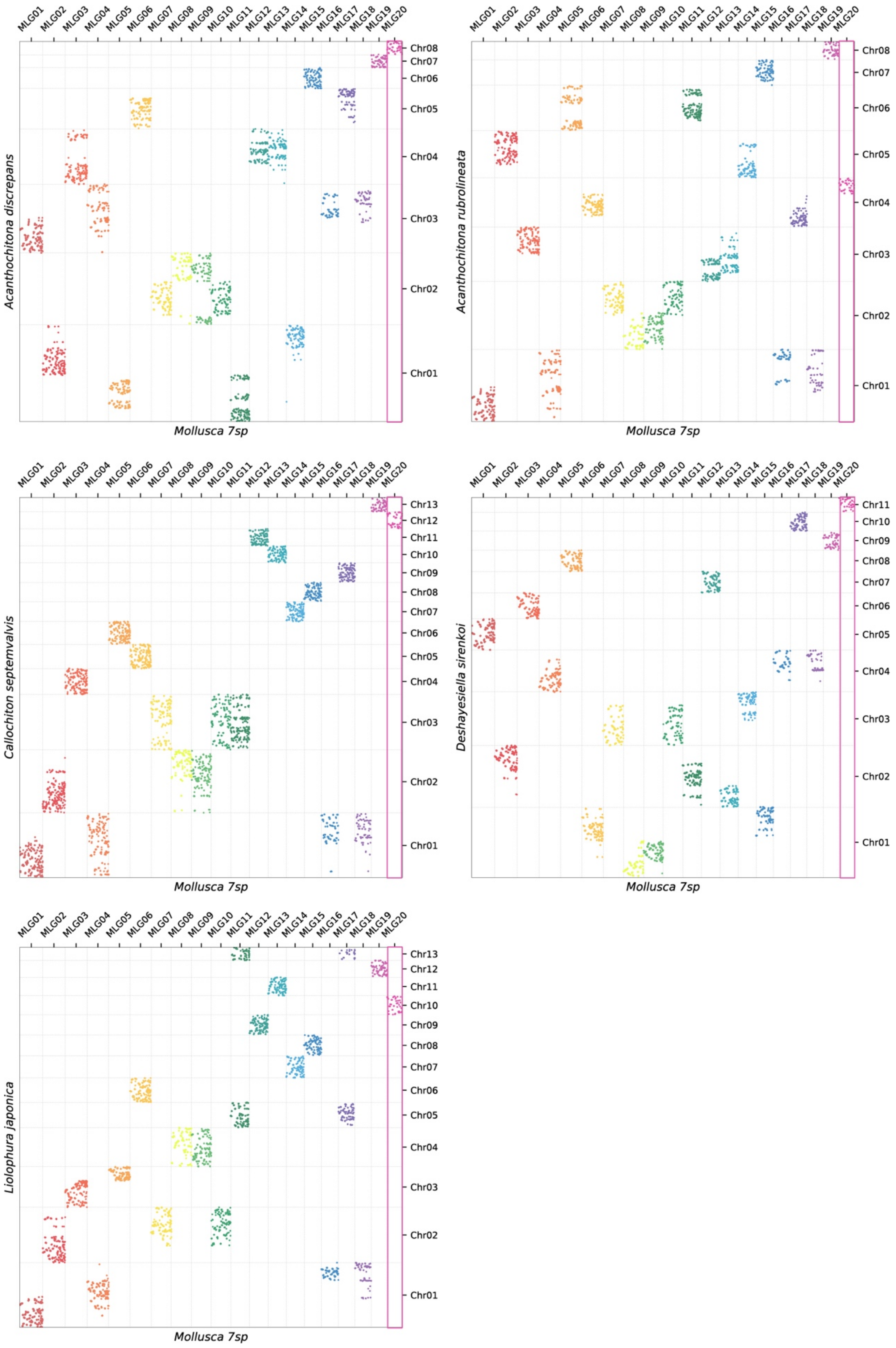


**Figure S4.** Oxford plots of 5 chitons against MLGs

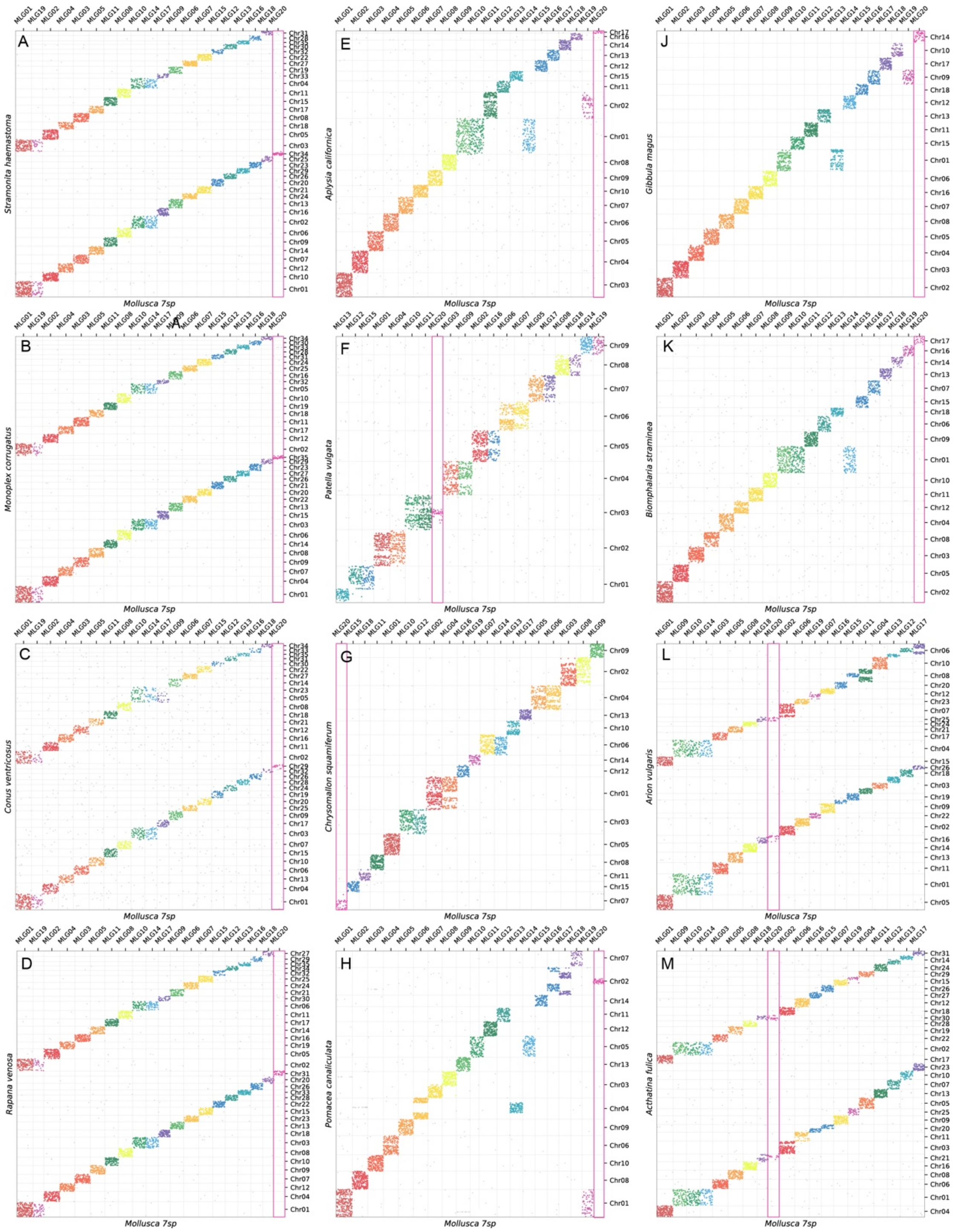


**Figure S5.** Oxford plots of 12 gastropods against MLGs, with the identification of the true sense of whole genome duplications (WGD). MLG20 is highlighted with a red box to distinguish its dynamics under WGD.

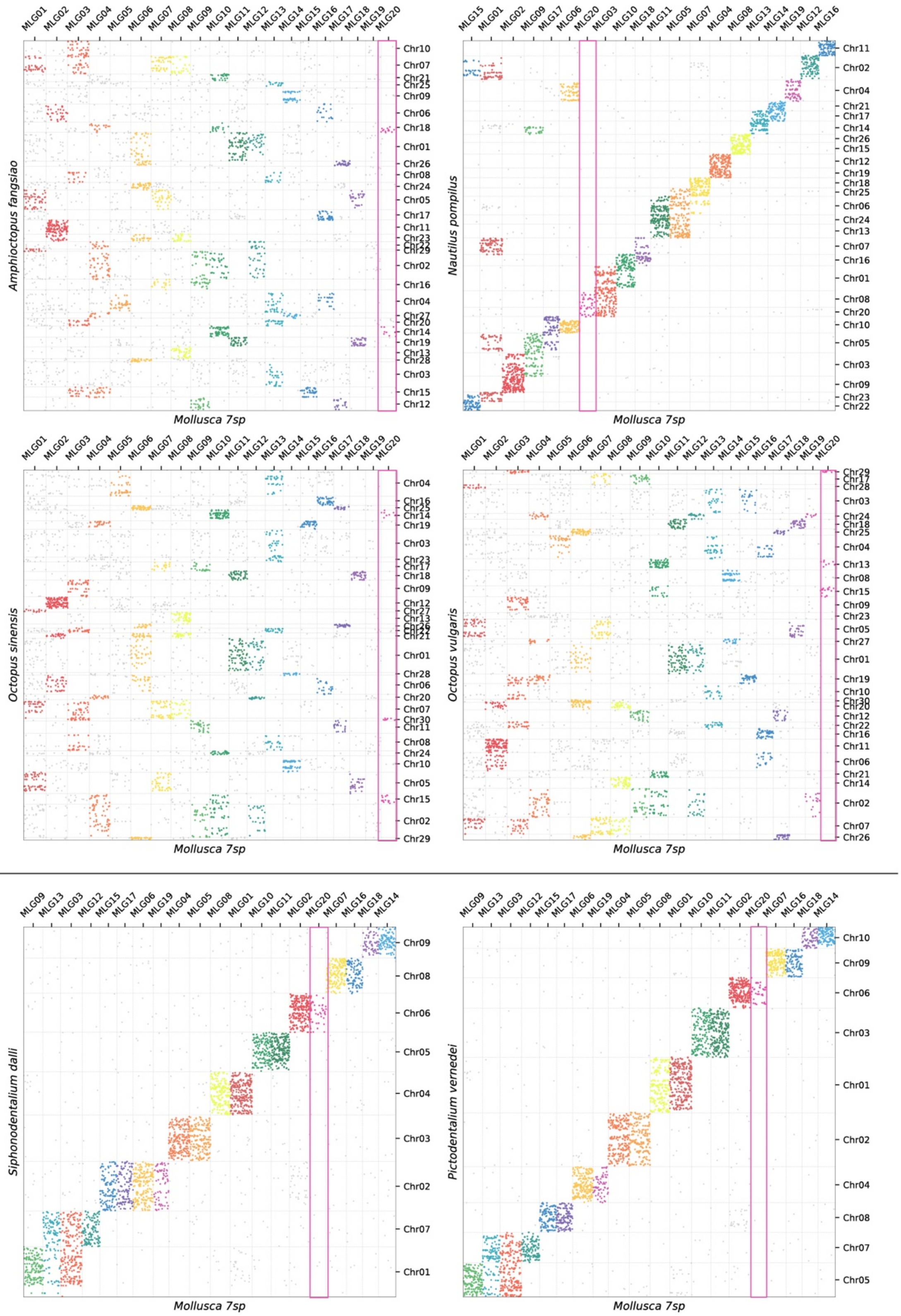


**Figure S6.** Oxford plots of 4 cephalopods and 2 scaphopods against MLGs.

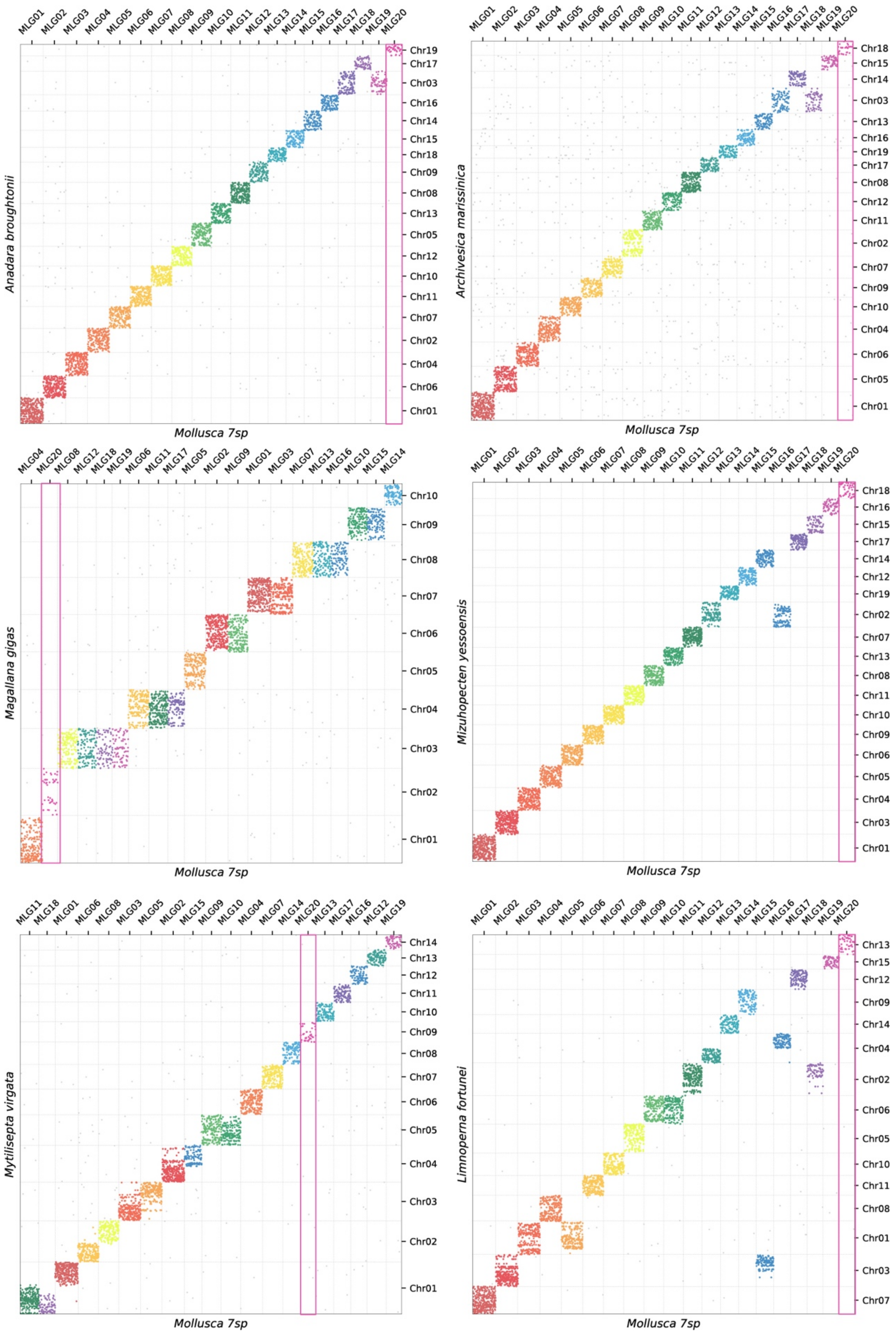


**Figure S7.** Oxford plots of 6 bivalves against MLGs.

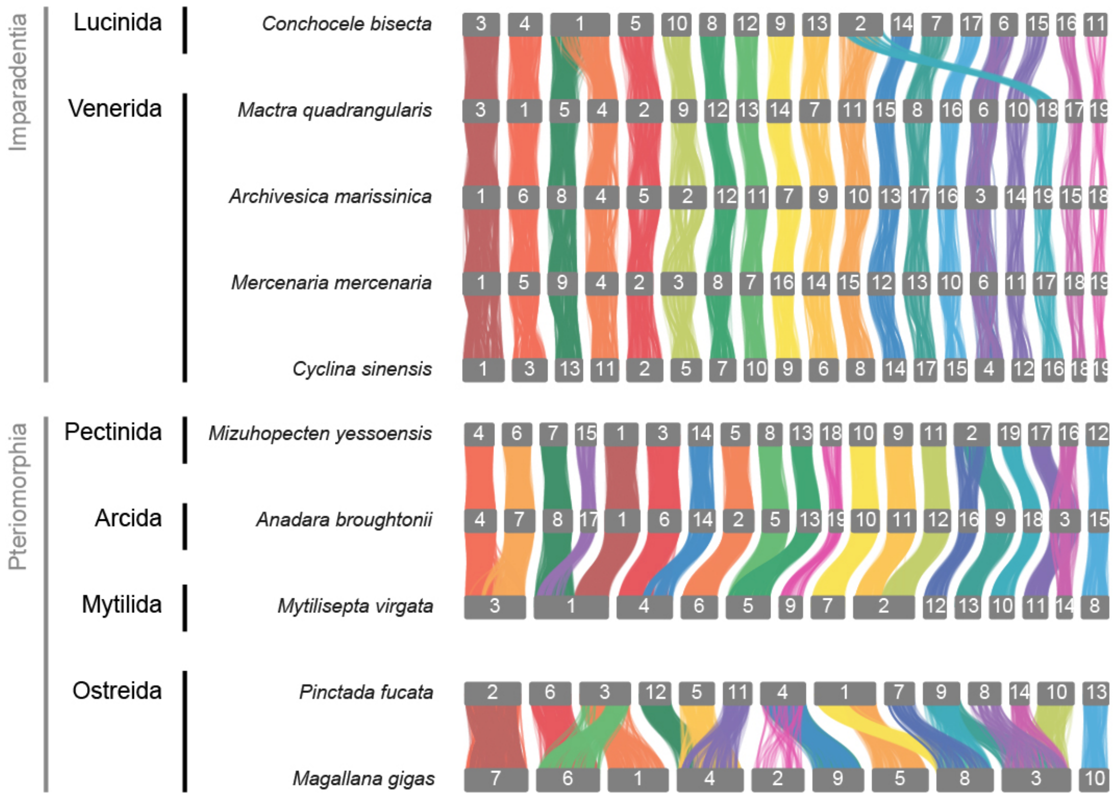


**Figure S8.** Syntenic graph in Bivalvia, highly conserved synteny in Imparadentia and large rearrangements within Peteriomorphia. Two clades in Bivalvia have 19 pseudo-chromosomes from a single fusion, but they are independent occurrences: in Pectinida, the fusion is MLG12 and MLG16, whereas in Venerida, the fusion is MLG16 and MLG18.

Supplementary tables to
Sigwart *et al.* Still waters run deep: Large scale genome rearrangement in the evolution of morphologically conservative Polyplacophora

**Table S1** The statistics of sequencing in four genomes, including HiFi, RNA, HiC
**Table S2** Genome assemblies and annotations
**Table S3** Comparison in chitons genomes
**Table S4** Nonsyntenic rate between species
**Table S5** Translocation rate at 25 species

**Table S1.** Summary statistics from genome sequencing of four new chiton genomes, including HiFi and Hi-C

| | | *Deshayesiella sirenkoi* | *Callochiton septemvalvis* | *Acanthochitona discrepans* | *Acanthochitona rubrolineata* |
|---|---|---|---|---|---|
| | Sampling area | Daikoku vent field, W Pacific Ocean | intertidal of Strangford Lough, N Ireland | | intertidal of Quingdao, China |
| HiFi | Number of reads | 4,713,356 | 8,086,367 | 5,785,025 | 3,675,326 |
| | Sum of reads | 78,716,116,715 | 39,763,935,831 | 30,682,344,156 | 57,655,991,192 |
| Hi-C (PE150) | Number of reads | 813,659,868 | 2,040,374,584 | 302,327,576 | 778,302,886 |
| | Sum of reads | 122,048,980,200 | 306,056,187,600 | 45,349,136,400 | 77,830,288,600 |
| RNA-seq (PE150) | Number of reads | 266,566,140 | 674,717,978 | 79,653,804 | 1,538,788,440 |
| | Sum of reads | 39,984,921,000 | 101,207,696,700 | 11,948,070,600 | 230,477,194,919 |

**Table S2.** Genome assemblies and gene-model prediction for four new chiton genomes.

| | | *Acanthochitona discrepans* | *Acanthochitona rubrolineata* |
|---|---|---|---|
| | Number | 704 | 165 |
| | Total base | 1,173,574,984 | 1,093,798,424 |
| contig-level | contig > 1M (number / total size) | | 52 / 1,075,127,699 |
| | L50 / N50 | 88 / 4,185,741 | 12 / 31,466,278 |
| | Completeness | C:97.4%[S:95.0%,D:2.4%],F:1.3%,M:1.3% | C:97.4%[S:95.0%,D:2.4%],F:1.8%,M:0.8%,n:954 |
| | Number | 396 | 112 |
| | Total base | 1,173,685,984 | 1,093,855,924 |
| seuchromosome-leve | Chromosomes | 8 | 8 |
| | Anchored rate | 94.34% | 98.39% |
| | Completeness | C:95.5%[S:93.9%,D:1.6%],F:1.0%,M:3.5%,n:954 | C:97.2%[S:95.3%,D:1.9%],F:2.0%,M:0.8%,n:954 |
| | Number of genes (longest isoform) | 21027 | 20,043 |
| | Completeness | C:95.5%[S:93.9%,D:1.6%],F:1.0%,M:3.5%,n:954 | C:95.6%[S:93.9%,D:1.7%],F:1.7%,M:2.7%,n:954 |
| Gene-model | Number of proteins (all isoforms) | 21027 | 30,176 |
| | Completeness | C:95.5%[S:93.9%,D:1.6%],F:1.0%,M:3.5%,n:954 | C:95.6%[S:75.9%,D:19.7%],F:1.7%,M:2.7%,n:954 |
| | Nr | 18,950 | 27,177 |
| | GO | 16,531 | 22,481 |
| | KEGG | 8,723 | 15,195 |

**Table S2.** (continued) Genome assemblies and gene-model prediction for four new chiton genomes.

| | | *Callochiton septemvalvis* | *Deshayesiella sirenkoi* |
|---|---|---|---|
| contig-level | Number | 3,543 | 358 |
| | Total base | 1,229,767,851 | 1,546,234,380 |
| | contig > 1M (number / total size) | 200 / 258,518,145 | 103 / 1,522,658,405 |
| | L50 / N50 | 663 / 619,644 | 19 / 27,005,026 |
| | Completeness | C:94.0%[S:90.9%,D:3.1%],F:3.0%,M:3.0%,n:954 | C:97.0%[S:96.2%,D:0.8%],F:2.0%,M:1.0%,n:954 |
| pseuchromosome-leve | Number | 416 | 207 |
| | Total base | 1,231,359,351 | 1,546,349,880 |
| | Chromosomes | 13 | 11 |
| | Anchored rate | 96.33% | 99.30% |
| | Completeness | C:94.2%[S:91.7%,D:2.5%],F:2.8%,M:3.0%,n:954 | C:97.0%[S:96.4%,D:0.6%],F:2.0%,M:1.0%,n:954 |
| Gene-model | Number of genes (longest isoform) | 19,591 | 23,050 |
| | Completeness | C:93.2%[S:90.0%,D:3.2%],F:2.2%,M:4.6%,n:954 | C:95.0%[S:94.1%,D:0.9%],F:1.2%,M:3.8%,n:954 |
| | Number of proteins (all isoforms) | 23,429 | 28,132 |
| | Completeness | C:93.3%[S:81.8%,D:11.5%],F:2.2%,M:4.5%,n:954 | C:95.1%[S:80.7%,D:14.4%],F:1.2%,M:3.7%,n:954 |
| | Nr | 21,256 | 25,312 |
| | GO | 17,456 | 16,145 |
| | KEGG | 11,313 | 11,232 |

**Table S3.** Comparison of summary data among available chitons genomes; the four new chiton genomes are indicated in **bold text**. N.B. analyses also incorporate the other chromosome-level genome for *Liolophura japonica*, which was published prior to the present study.

| Order: Family | Species | Reference | Assembly level | Chromosomes | Genome size (bp) | N50 (bp) / L50 |
|---|---|---|---|---|---|---|
| **Lepidopleurida: Protochitonidae** | ***Deshayesiella sirenkoi*** | **this study** | **chromosome** | **11** | **1,546,349,880** | **169,292,669 / 4** |
| Lepidopleurida: Hanleyidae | *Hanleya hanleyi* | Varney et al., 2022 | contig | unknown | 2,516,608,230 | 65,037 / 10427 |
| **Callochitonida: Callochitonidae** | ***Callochiton septemvalvis*** | **this study** | **chromosome** | **13** | **1,231,359,351** | **80,830,436 / 4** |
| Chitonida: Schizochitonidae | *Schizochiton incissus* | Liu et al., 2023 | contig | unknown | 585,667,302 | 25,469 / 6,633 |
| Chitonida: Chitonidae | *Liolophura japonica* | HKBGC, 2024 | chromosome | 13 | 609,495,693 | 37,343,639 |
| Chitonida: Chitonidae | *Acanthopleura granulata* | Varney et al., 2020 | contig | unknown | 606,536,932 | 23,921,462 / 9 |
| **Chitonida: Acanthochitonidae** | ***Acanthochitona rubrolineata*** | **this study** | **chromosome** | **8** | **1,093,855,924** | **154,826,102 / 3** |
| **Chitonida: Acanthochitonidae** | ***Acanthochitona discrepans*** | **this study** | **chromosome** | **8** | **1,173,685,984** | **199,945,981 / 3** |

**Table S3. (continued)** Comparison of summary data among available chitons genomes; the four new chiton genomes are indicated in **bold text**. N.B. analyses also incorporate the other chromosome-level genome for *Liolophura japonica,* which was published prior to the

| Completeness (genome, busco 5.2.2, metazoa_odb10) | Contigs or Scaffolds | Completeness (the longest coding isoforms) | Number of proteins (the longest isoforms) |
|---|---|---|---|
| **C:97.0%[S:96.2%,D:0.8%],F:2.0%,M:1.0%** | **207** | **C:95.0%[S:94.1%,D:0.9%],F:1.2%,M:3.8%** | **23,050** |
| C:79.4%[S:74.4%,D:5.0%],F:12.6%,M:8.0% | 57,495 | C:81.8%[S:75.6%,D:6.2%],F:11.1%,M:7.1% | 69,284 |
| **C:94.0%[S:90.9%,D:3.1%],F:3.0%,M:3.0%** | **416** | **C:93.2%[S:90.0%,D:3.2%],F:2.2%,M:4.6%** | **19,591** |
| C:63.5%[S:58.2%,D:5.3%],F:14.5%,M:22.0% | 25,461 | C:40.9%[S:37.5%,D:3.4%],F:19.8%,M:39.3% | 20,902 |
| C:96.2%[S:95.6%,D:0.6%],F:2.0%,M:1.8% | 632 | C:91.0%[S:90.3%,D:0.7%],F:4.9%,M:4.1% | 25,163 |
| C:95.8%[S:95.2%,D:0.6%],F:2.3%,M:1.9% | 87 | C:94.0%[S:93.4%,D:0.6%],F:2.4%,M:3.6% | 20,470 |
| **C:97.4%[S:95.0%,D:2.4%],F:1.8%,M:0.8%** | **112** | **C:95.6%[S:93.9%,D:1.7%],F:1.7%,M:2.7%** | **20,043** |
| **C:97.4%[S:95.0%,D:2.4%],F:1.3%,M:1.3%** | **396** | **C:95.5%[S:93.9%,D:1.6%],F:1.0%,M:3.5%** | **21,027** |

**Table S4.** Nonsyntenic rate between species (summarised by clade in Table S5) following the method described in the Supporting Information appendix.

| | *Conchocele bisecta* | *Mactra quadrangularis* | *Mizuhopecten yessoensis* | *Limnoperna fortunei* | *Mytilisepta virgata* | *Mytilus unguiculatus* | *Pinctada fucata* | *Magallana gigas* |
|---|---|---|---|---|---|---|---|---|
| *Conchocele bisecta* | | | | | | | | |
| *Mactra quadrangularis* | 12.29 | | | | | | | |
| *Mizuhopecten yessoensis* | 12.37 | 14.35 | | | | | | |
| *Limnoperna fortunei* | 12.02 | 14.70 | 9.42 | | | | | |
| *Mytilisepta virgata* | 12.86 | 15.12 | 10.30 | 6.60 | | | | |
| *Mytilus unguiculatus* | 14.99 | 16.63 | 12.20 | 9.03 | 9.85 | | | |
| *Pinctada fucata* | 13.89 | 16.65 | 11.91 | 12.57 | 13.32 | 15.23 | | |
| *Magallana gigas* | 13.16 | 15.87 | 11.41 | 11.86 | 11.91 | 13.65 | 12.94 | |
| *Patella vulgata* | 13.43 | 15.47 | 12.32 | 12.13 | 12.34 | 13.51 | 13.81 | 12.77 |
| *Gibbula magus* | 12.16 | 14.82 | 10.85 | 11.16 | 11.80 | 13.96 | 13.02 | 12.51 |
| *Chrysomallon squamiferum* | 10.60 | 12.67 | 9.25 | 9.93 | 9.94 | 10.72 | 10.88 | 10.73 |
| *Gigantopelta aegis* | 11.76 | 13.63 | 9.88 | 10.33 | 10.53 | 12.15 | 11.63 | 10.85 |
| *Aplysia californica* | 13.71 | 15.49 | 12.66 | 12.94 | 13.52 | 14.46 | 14.15 | 13.51 |
| *Biomphalaria straminea* | 14.23 | 15.68 | 11.99 | 11.80 | 13.38 | 14.56 | 13.21 | 12.82 |
| *Arion vulgaris* | 13.81 | 14.87 | 12.55 | 12.63 | 12.72 | 14.36 | 13.72 | 12.81 |
| *Acthatina fulica* | 14.38 | 15.90 | 12.99 | 13.30 | 15.68 | 16.47 | 14.36 | 13.66 |
| *Pomacea canaliculata* | 12.26 | 14.10 | 10.68 | 11.27 | 11.23 | 12.78 | 12.95 | 11.55 |
| *Monoplex corrugatus* | 13.14 | 15.67 | 11.66 | 11.64 | 12.21 | 13.09 | 12.87 | 12.37 |
| *Deshayesiella sirenkoi* | 11.59 | 13.56 | 10.04 | 9.97 | 10.13 | 11.35 | 11.17 | 10.38 |
| *Callochiton septemvalvis* | 18.70 | 20.22 | 17.36 | 16.91 | 16.84 | 17.88 | 18.07 | 16.93 |
| *Acanthochitona discrepans* | 10.88 | 13.00 | 10.29 | 10.01 | 10.27 | 10.99 | 11.09 | 9.56 |
| *Acanthochitona rubrolineata* | 11.14 | 12.94 | 9.90 | 9.52 | 10.42 | 10.92 | 10.93 | 9.88 |
| *Liolophura japonica* | 13.07 | 15.02 | 11.13 | 11.27 | 12.04 | 13.21 | 12.95 | 11.15 |
| *Pictodentalium vernedei* | 14.19 | 16.64 | 12.78 | 12.37 | 13.36 | 14.52 | 13.24 | 12.22 |
| *Siphonodentalium dalli* | 17.41 | 19.28 | 15.58 | 15.38 | 14.74 | 17.17 | 15.58 | 14.29 |

**Table S4. (continued)** Nonsyntenic rate between species (summarised by clade in Table S5) following the method described in the Supporting Information appendix.

| | *Patella vulgata* | *Gibbula magus* | *Chrysomallon squamiferum* | *Gigantopelta aegis* | *Aplysia californica* | *Biomphalaria straminea* | *Arion vulgaris* | *Acthatina fulica* |
|---|---|---|---|---|---|---|---|---|
| *Conchocele bisecta* | | | | | | | | |
| *Mactra quadrangularis* | | | | | | | | |
| *Mizuhopecten yessoensis* | | | | | | | | |
| *Limnoperna fortunei* | | | | | | | | |
| *Mytilisepta virgata* | | | | | | | | |
| *Mytilus unguiculatus* | | | | | | | | |
| *Pinctada fucata* | | | | | | | | |
| *Magallana gigas* | | | | | | | | |
| *Patella vulgata* | | | | | | | | |
| *Gibbula magus* | 10.25 | | | | | | | |
| *Chrysomallon squamiferum* | 9.50 | 6.63 | | | | | | |
| *Gigantopelta aegis* | 10.08 | 7.57 | 2.93 | | | | | |
| *Aplysia californica* | 13.75 | 11.26 | 11.08 | 11.25 | | | | |
| *Biomphalaria straminea* | 12.28 | 10.40 | 10.09 | 10.40 | 7.28 | | | |
| *Arion vulgaris* | 13.19 | 9.67 | 10.76 | 11.83 | 5.76 | 5.55 | | |
| *Acthatina fulica* | 15.00 | 11.47 | 11.48 | 12.30 | 7.52 | 7.43 | 11.14 | |
| *Pomacea canaliculata* | 11.02 | 8.56 | 8.98 | 9.26 | 12.07 | 11.03 | 11.30 | 12.23 |
| *Monoplex corrugatus* | 10.71 | 8.72 | 7.84 | 9.69 | 12.96 | 13.86 | 20.21 | 23.23 |
| *Deshayesiella sirenkoi* | 9.66 | 9.24 | 8.32 | 8.52 | 11.08 | 9.91 | 10.97 | 12.27 |
| *Callochiton septemvalvis* | 17.38 | 16.76 | 15.22 | 15.81 | 17.80 | 18.22 | 18.42 | 19.62 |
| *Acanthochitona discrepans* | 9.34 | 9.28 | 8.01 | 8.13 | 11.18 | 10.11 | 10.55 | 11.40 |
| *Acanthochitona rubrolineata* | 9.74 | 8.96 | 8.33 | 8.36 | 11.32 | 10.03 | 10.91 | 12.52 |
| *Liolophura japonica* | 11.60 | 10.50 | 9.81 | 9.89 | 12.83 | 11.37 | 12.96 | 13.62 |
| *Pictodentalium vernedei* | 13.55 | 11.93 | 10.79 | 12.40 | 14.93 | 12.82 | 15.96 | 17.61 |
| *Siphonodentalium dalli* | 15.39 | 15.28 | 13.35 | 14.41 | 17.58 | 16.84 | 19.64 | 20.76 |

**Table S4. (continued)** Nonsyntenic rate between species (summarised by clade in Table S5) following the method described in the Supporting Information appendix.

| | *Pomacea canaliculata* | *Monoplex corrugatus* | *Deshayesiella sirenkoi* | *Callochiton septemvalvis* | *Acanthochiton a discrepans* | *Acanthochitona rubrolineata* | *Liolophura japonica* | *Pictodentalium vernedei* |
|---|---|---|---|---|---|---|---|---|
| *Conchocele bisecta* | | | | | | | | |
| *Mactra quadrangularis* | | | | | | | | |
| *Mizuhopecten yessoensis* | | | | | | | | |
| *Limnoperna fortunei* | | | | | | | | |
| *Mytilisepta virgata* | | | | | | | | |
| *Mytilus unguiculatus* | | | | | | | | |
| *Pinctada fucata* | | | | | | | | |
| *Magallana gigas* | | | | | | | | |
| *Patella vulgata* | | | | | | | | |
| *Gibbula magus* | | | | | | | | |
| *Chrysomallon squamiferum* | | | | | | | | |
| *Gigantopelta aegis* | | | | | | | | |
| *Aplysia californica* | | | | | | | | |
| *Biomphalaria straminea* | | | | | | | | |
| *Arion vulgaris* | | | | | | | | |
| *Acthatina fulica* | | | | | | | | |
| *Pomacea canaliculata* | | | | | | | | |
| *Monoplex corrugatus* | 6.75 | | | | | | | |
| *Deshayesiella sirenkoi* | 9.02 | 11.17 | | | | | | |
| *Callochiton septemvalvis* | 16.57 | 20.22 | 12.35 | | | | | |
| *Acanthochitona discrepans* | 9.64 | 10.99 | 4.65 | 11.62 | | | | |
| *Acanthochitona rubrolineata* | 9.45 | 11.11 | 4.13 | 11.66 | 2.51 | | | |
| *Liolophura japonica* | 10.91 | 12.58 | 4.69 | 12.09 | 3.85 | 3.69 | | |
| *Pictodentalium vernedei* | 11.73 | 14.22 | 10.57 | 16.71 | 8.61 | 9.11 | 10.78 | |
| *Siphonodentalium dalli* | 13.60 | 20.32 | 12.31 | 17.98 | 11.50 | 11.36 | 12.97 | 13.56 |

**Table S5.** Summary information for translocation rate of 25 species of molluscs, organised by taxonomic class.

| Class | Species | Chromosomes | Nonsyntenic rate (%) | Translocation rate ( Mya per 1%) | Mean (Class) |
|---|---|---|---|---|---|
| Bivalvia | *Conchocele bisecta* | 17 | 13.32 | 39.79 | 41.10 |
| | *Mactra quadrangularis* | 19 | 15.23 | 34.79 | |
| | *Mizuhopecten yessoensis* | 19 | 11.88 | 44.62 | |
| | *Limnoperna fortunei* | 15 | 11.91 | 44.49 | |
| | *Mytilisepta virgata* | 14 | 12.42 | 42.67 | |
| | *Mytilus unguiculatus* | 14 | 13.65 | 38.82 | |
| | *Pinctada fucata* | 14 | 13.16 | 40.29 | |
| | *Magallana gigas* | 14 | 12.23 | 43.32 | |
| Gastropoda | *Patella vulgata* | 9 | 12.83 | 41.31 | 41.87 |
| | *Gibbula magus* | 18 | 12.15 | 43.63 | |
| | *Chrysomallon squamiferum* | 15 | 10.57 | 50.15 | |
| | *Gigantopelta aegis* | 14 | 11.22 | 47.25 | |
| | *Aplysia californica* | 17 | 13.81 | 38.38 | |
| | *Biomphalaria straminea* | 18 | 13.13 | 40.36 | |
| | *Arion vulgaris* | 26 | 13.79 | 38.43 | |
| | *Acthatina fulica* | 31 | 14.97 | 35.40 | |
| | *Pomacea canaliculata* | 14 | 11.85 | 44.73 | |
| | *Monoplex corrugatus* | 35 | 13.55 | 39.11 | |
| Polyplacophora | *Deshayesiella sirenkoi* | 11 | 10.56 | 50.18 | 45.48 |
| | *Callochiton septemvalvis* | 13 | 17.68 | 29.98 | |
| | *Acanthochitona discrepans* | 8 | 10.24 | 51.76 | |
| | *Acanthochitona rubrolineata* | 8 | 10.34 | 51.25 | |
| | *Liolophura japonica* | 13 | 11.98 | 44.23 | |
| Scaphopoda | *Pictodentalium vernedei* | 10 | 13.09 | 40.50 | 37.05 |
| | *Siphonodentalium dalli* | 9 | 15.77 | 33.61 | |
